# Noncanonical structural requirements of neurofibromin SUMOylation reveal a folding-deficiency of several pathogenic mutants

**DOI:** 10.1101/2021.12.09.471973

**Authors:** Mohammed Bergoug, Christine Mosrin, Fabienne Godin, Michel Doudeau, Iva Sosic, Marcin Suskiewicz, Béatrice Vallée, Hélène Bénédetti

## Abstract

Neurofibromin (Nf1) is a large multidomain protein encoded by the tumour-suppressor gene *NF1*. *NF1* is mutated in a frequently occurring genetic disease, neurofibromatosis type I, and in various cancers. The best described function of Nf1 is its Ras-GTPase activity, carried out by its GAP-related domain (GRD). SecPH, another structurally well-characterized domain of Nf1, is immediately adjacent to the GRD and interacts with lipids and proteins, thus connecting Nf1 to diverse signalling pathways. Here, we demonstrate, for the first time, that Nf1 and SecPH are substrates of the SUMO pathway. We identified a well-defined SUMOylation profile of SecPH and a main SUMOylation event on Lys1731 that appears to play a role in Ras-GAP activity. Our data allowed us to characterize a new set of pathogenic Nf1 missense mutants that exhibits a disrupted SUMOylation profile that may correlate with their unfolding. Accordingly, Lys1731 SUMOylation is mediated by a noncanonical structural motif, therefore allowing a read-out of SecPH conformation and folding status.

## INTRODUCTION

Neurofibromin is encoded by the *NF1* gene, which is responsible for neurofibromatosis type I (NF1), a genetic disease that affects approximately one individual in 3,500. The disease is characterized by a wide spectrum of manifestations: the development of various types of peripheral nervous system and brain tumours, pigmentary lesions, skeletal abnormalities, learning disabilities, attention deficits, and social and behavioural problems (for reviews see^1,2,3,4^). *NF1* is also mutated in various types of cancer^5, 6,7,8,9^. More than 3,000 different germline *NF1* mutations have been reported in the Human Gene Mutation Database, revealing a substantial diversity of pathogenetic mutations with no single hotspot^10^.

Nf1 is a large dimeric^11^ multidomain protein of 2,818 residues (for a review of Nf1 structure, function, and regulation, see ^12^). It contains an N-terminal cysteine-serine rich domain (CSRD) and a central GAP (GTPase-activating protein)-related domain (GRD), followed by a SecPH domain, a bipartite module composed of a segment homologous to the yeast Sec-14 phospholipid interaction module and a pleckstrin homology (PH)-like domain, and, finally, a C-terminal domain (CTD). The 3D structures of GRD and SecPH domains have long been the only one to be resolved ^13,14^, but recently, the 3D structure of the entire Nf1 dimer has been elucidated by cryo EM^15^.

The best characterized function of Nf1 is its Ras-GAP activity, which is carried out by its GRD. This domain increases the intrinsic GTPase activity of Ras by a transition state-stabilizing mechanism^13, 16^. Nf1 also regulates cAMP levels in various cell types in both a Ras-dependent and Ras-independent manner^17, 18^, thus influencing tumour formation, as well as neurite growth and the formation of growth cones into neurons of the CNS. Finally, Nf1 plays an important role in regulating actin cytoskeletal dynamics through the negative regulation of the Rho/ROCK/LIMK2/cofilin^19^ and Rac1/Pak1/LIMK1/cofilin pathways^20^.

Due to its proximity to the GRD and its ability to bind phospholipids^14, 21^ and proteins^19^, SecPH has been hypothesized to allosterically regulate the Ras-GAP activity of the GRD. However, the role played by SecPH/lipid interactions in Nf1 function is still unknown. By contrast, the ability of SecPH to mediate interactions with actors of new signalling pathways (LIMK2, R5HT6) led to the identification of new Nf1 functions^19, 22^.

The functions of Nf1 have been shown to be regulated by various post-translational modifications (PTMs), such as phosphorylation, which can allosterically affect its Ras-GAP activity^23, 24^ and localization^25^, and ubiquitylation, which affects its stability^26^.

SUMOylation is a PTM that, similarly to <u>ubiquitination</u>, involves the covalent enzymatic conjugation of the SUMO (Small Ubiquitin-like Modifier) protein to specific lysine residues of substrate proteins^27, 28^. It involves a dedicated enzymatic pathway, comprising the heterodimeric E1-activating enzyme SAE1-SAE2, the sole E2 conjugating enzyme Ubc9, and various E3 ligases, which can be optional^29^. SUMOylation is a reversible process due to the activity of specific deSUMOylation enzymes called sentrin-proteases (SENP)^30^. At the molecular level, SUMOylation can modulate the activity, stability, localization, and interaction pattern of target proteins^31^.

The SUMOylation of Nf1 was first suggested by our group, which showed partial colocalization of Nf1 with the Promyelocytic Leukemia Protein (PML) in PML nuclear bodies, dynamic multiprotein complexes that contain the SUMOylation machinery and suspected to be hotspots of SUMOylation^32^. Mass spectrometry-based global studies of SUMOylation have identified Nf1 as a potential SUMO target protein^33^ and two SUMOylated lysines belonging to the 23a alternative exon, located in the Nf1 GRD, were identified^34^. However, the SUMOylation of Nf1 has never been further characterized.

Here, we demonstrate that Nf1 and its SecPH domain are SUMOylated by the SUMO2 paralog. SecPH exhibited a specific SUMOylation profile involving the major SUMOylation of a specific lysine, K1731. This major SUMOylation had a functional impact on Nf1 function by stimulating Nf1 Ras-GAP. Furthermore, five of the nine *NF1* pathogenic variants that we studied showed a well-defined modified SUMOylation pattern, lacking K1731 SUMOylation and, instead, exhibiting high molecular-weight species and instability suggestive of a folding deficiency. Surprisingly, K1731 SUMOylation was independent of the linear SUMOylation consensus sequence flanking this site, instead being mediated by a noncanonical structural motif. Overall, our results suggest that SecPH SUMOylation could serve as a readout of its folding status and conformation to further induce its proteasomal degradation or regulate a number of its functions.

## RESULTS

### 1. Endogenous Nf1 is preferentially SUMOylated by SUMO2

We first validated Nf1 as a true SUMO substrate by co-transfecting HEK293 cells, which naturally produce Nf1, with two plasmids allowing the production of the SUMO-conjugating enzyme, Ubc9, and Flag-tagged SUMO1 or SUMO2 (which should also behave as SUMO3, as they exhibit 97% peptide sequence similarity). Mock-transfected cells were used as a negative control. Non-transfected and transfected cells showed equal Nf1 levels by western blotting (Fig. 1a) and Flag-SUMO1 and SUMO2 (15 kDa) were well expressed and differentially integrated into SUMOylated proteins, producing different patterns (Fig. 1b).

**Fig. 1.**
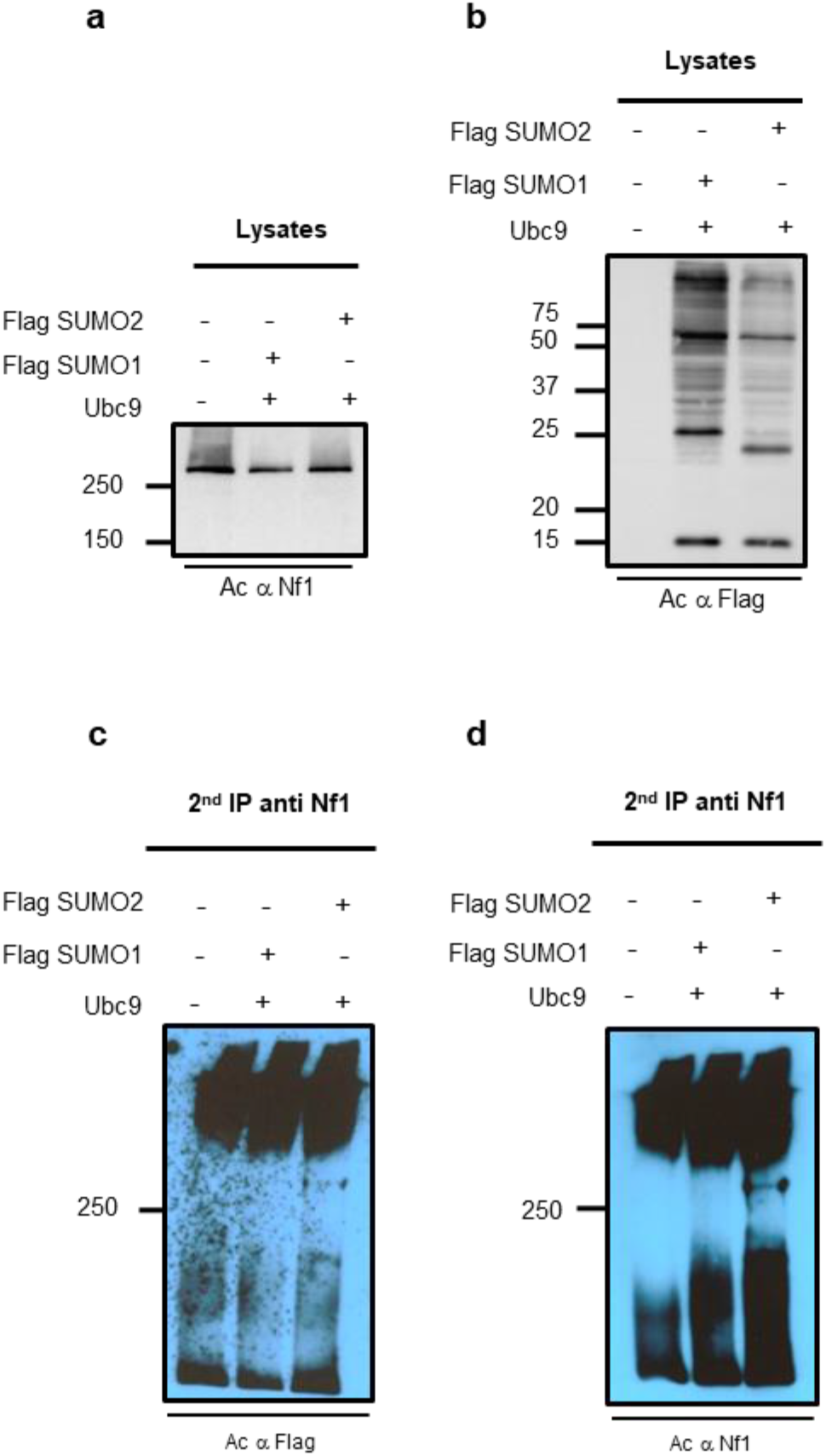
Endogenous Nf1 is SUMOylated. HEK293 cells were co-transfected with p3XFlag SUMO1 or p3XFlag SUMO2, along with pcDNA3 Ubc9. The corresponding empty plasmids were used in the transfection experiment as a negative control. Lysates were subjected to initial IP with anti-Flag antibodies for the enrichment of SUMOylated proteins or SUMO-interacting proteins, followed by a second IP with anti-Nf1 antibodies. **a** and **b,** Western-blot analysis of total lysates with (**a**) anti-Nf1 and (**b**) anti-Flag antibodies. **c** and **d,** Western-blot analysis of the eluates of the second IP with (**c**) anti-Flag and (**d**) anti-Nf1 antibodies.

Then, we first immunoprecipitated cell lysates with anti-Flag antibodies to enrich them for SUMOylated or SUMO-interacting proteins. We then performed a second immunoprecipitation using anti-Nf1 antibodies on the eluates of the first immunoprecipitate to enrich them for Nf1-derived proteins. The second immunoprecipitation eluate was analysed by western blotting with anti-Flag and anti-Nf1 antibodies (Fig. 1c, d). A band migrating at a molecular mass of approximately 300 kDa, the size corresponding to Nf1, was revealed by both antibodies only in the presence of Flag-SUMO2 (Fig. 1c and 1d). The detection of Nf1 with the anti-Flag antibodies demonstrates that it was conjugated to Flag-SUMO2 and not simply interacting with SUMO2-conjugated proteins.

### 2. The SecPH domain of Nf1 is SUMOylated

Nf1 is a large multidomain protein of 2,818 residues. Two SUMOylation prediction software programs, SUMOPLOT (https://www.abgent.com/sumoplot) and JASSA^35^, were used to analyse its sequence, resulting in the prediction of 13 SUMOylation consensus (ΨKxE/D)^36^ or inverted consensus (E/DxKΨ)^37^ sites with a high score for at least one software program (Fig. 2a).

**Fig. 2.**
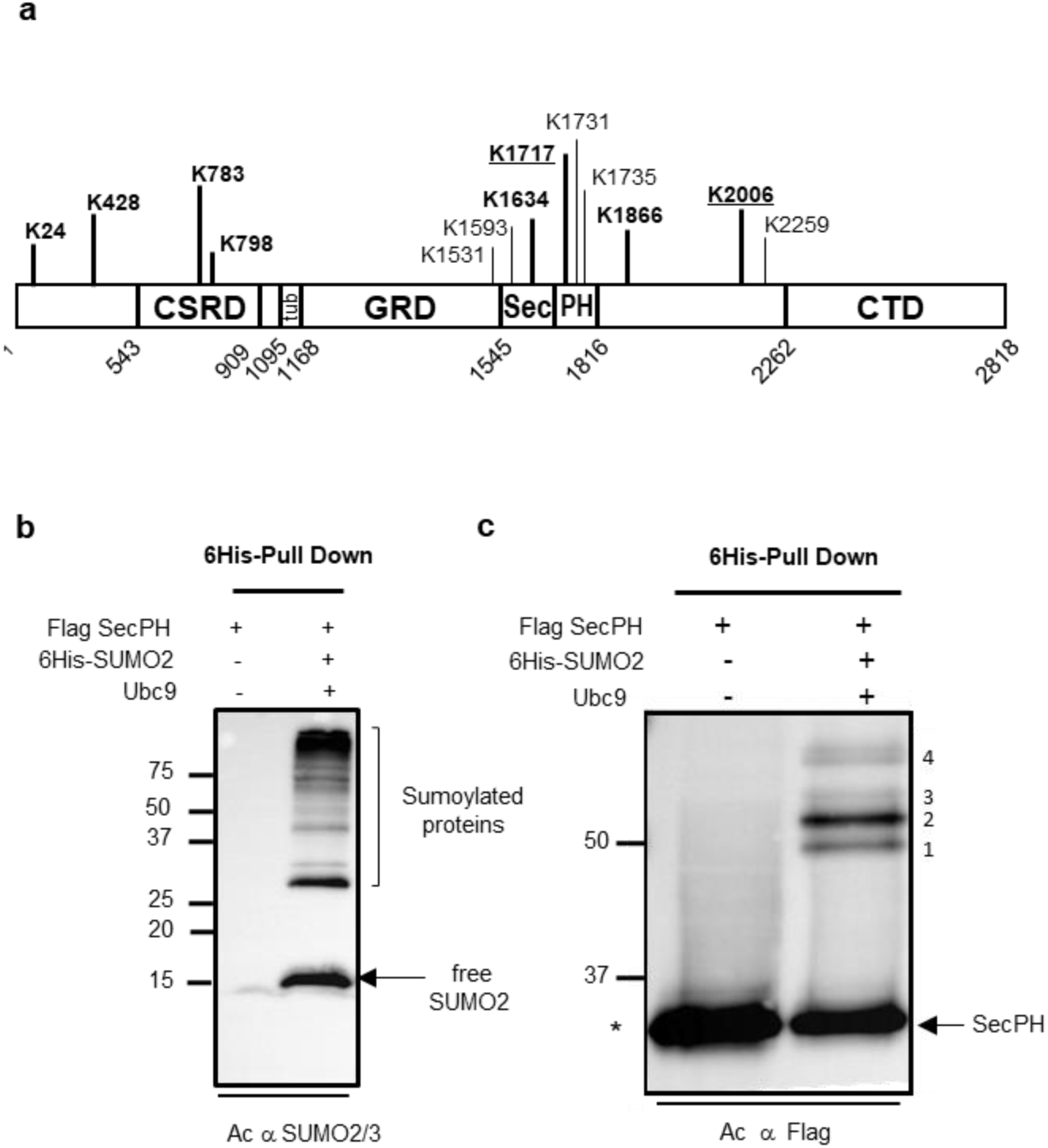
Localization of putative SUMO acceptor lysines in Nf1 domains and the presence of different forms of SUMOylation for the Nf1 SecPH domain. **a** SUMOylation consensus sites were predicted from analysis of the Nf1 sequence (NP_000258.1 neurofibromin isoform1 [Homo sapiens] 2818 aa protein) using the SUMOPLOT and JASSA bioinformatic tools. Lysines predicted to have a high probability of SUMOylation by both software programs are indicated in bold letters and underlined (**<u>K1717</u>**, **<u>K2006</u>**). Lysines predicted to have a high probability of SUMOylation by one software and a low probability by the other are indicated in bold (**K24, K428, K783, K798**, **K1634, K1866**). The other lysines were predicted to be SUMOylated with a high probability by only one software (K1531, K1593, K1731, K1735, K2259). **b** and **c,** HEK293 cells were co-transfected with p3XFlag SecPH, pcDNA3 6His-SUMO2, and pcDNA3 Ubc9 or only p3XFlag SecPH as a negative control. After 24 h of growth, cells were lysed in 6M guanidinium buffer and SUMOylated proteins enriched by pull-down using cobalt beads, as described in Methods. Pull-down eluates were analysed by western-blotting and revealed with (**b**) anti-SUMO2/3 or (**c**) anti-Flag antibodies. The four bands denoted 1 to 4 show the SUMOylated SecPH species. The asterisk indicates nonspecific binding of unmodified SecPH to the cobalt beads.

We decided to focus on the SecPH domain of Nf1 to further characterize its SUMOylation for several reasons : (i) the SecPH domain contains the highest number of SUMOylation consensus sites, (ii) its 3D structure has been studied in details^14, 21, 38^, (iii) it is adjacent to, and may regulate, the GRD domain, which carries the Ras-GAP activity of Nf1, and (iv) pleckstrin homology domains are enriched in SUMOylation substrates^39^.

We first studied SecPH SUMOylation by co-transfecting HEK293 cells with three plasmids to overproduce Flag-tagged SecPH, Ubc9, and 6His-tagged SUMO2. After lysis in denaturing conditions, SUMOylated proteins were enriched by performing a pull-down on cobalt beads. Western-blot analysis of eluates using anti-SUMO2/3 antibodies showed the 6His pull-down to be strongly enriched in SUMOylated proteins (Fig. 2b).

Eluates were also analysed by western blotting using anti-Flag antibodies. SUMOylated Flag-SecPH showed a very reproducible and discrete profile of four bands: three (bands 1, 2, and 3) migrated at approximately 50 kDa, whereas band 4 migrated with a slightly higher molecular weight (Fig. 2c).

### 3. K1634 and K1731 play a role in the SecPH SUMOylation profile

We next identified the SUMO conjugation sites within SecPH. Lysine residues belonging to bioinformatically-predicted putative SUMOylation sites were independently mutated into arginine, creating five SecPH mutants: SecPH K1593R, SecPH K1634R, SecPH K1717R, SecPH K1731R, and SecPH K1735R (Fig. 3a). The strongest SUMOylation band, band 2, disappeared from SecPH K1731R, for which the three remaining SUMOylation bands became slightly stronger relative to WT SecPH (Fig. 3b). A number of mutants exhibited a stronger signal than WT SecPH, especially SecPH K1634R. In this latter mutant, the resulting broader band 2 could have hidden changes in band 3. Thus, we constructed a SecPH K1634,K1731R double mutant and compared its SUMOylation profile to that of each corresponding SecPH single mutant. The western blot showed a supplementary band (called x) occurring just above band 1 in SecPH K1731R and SecPH K1634,K1731R and band 2 and 3 were absent from SecPH K1634,K1731R (Supplementary Fig. 1). These results indicate that K1731 and K1634 were required for the presence of bands 2 and 3, respectively, and that band x may have been hidden by band 2 or reflect a compensation phenomenon.

**Fig. 3.**
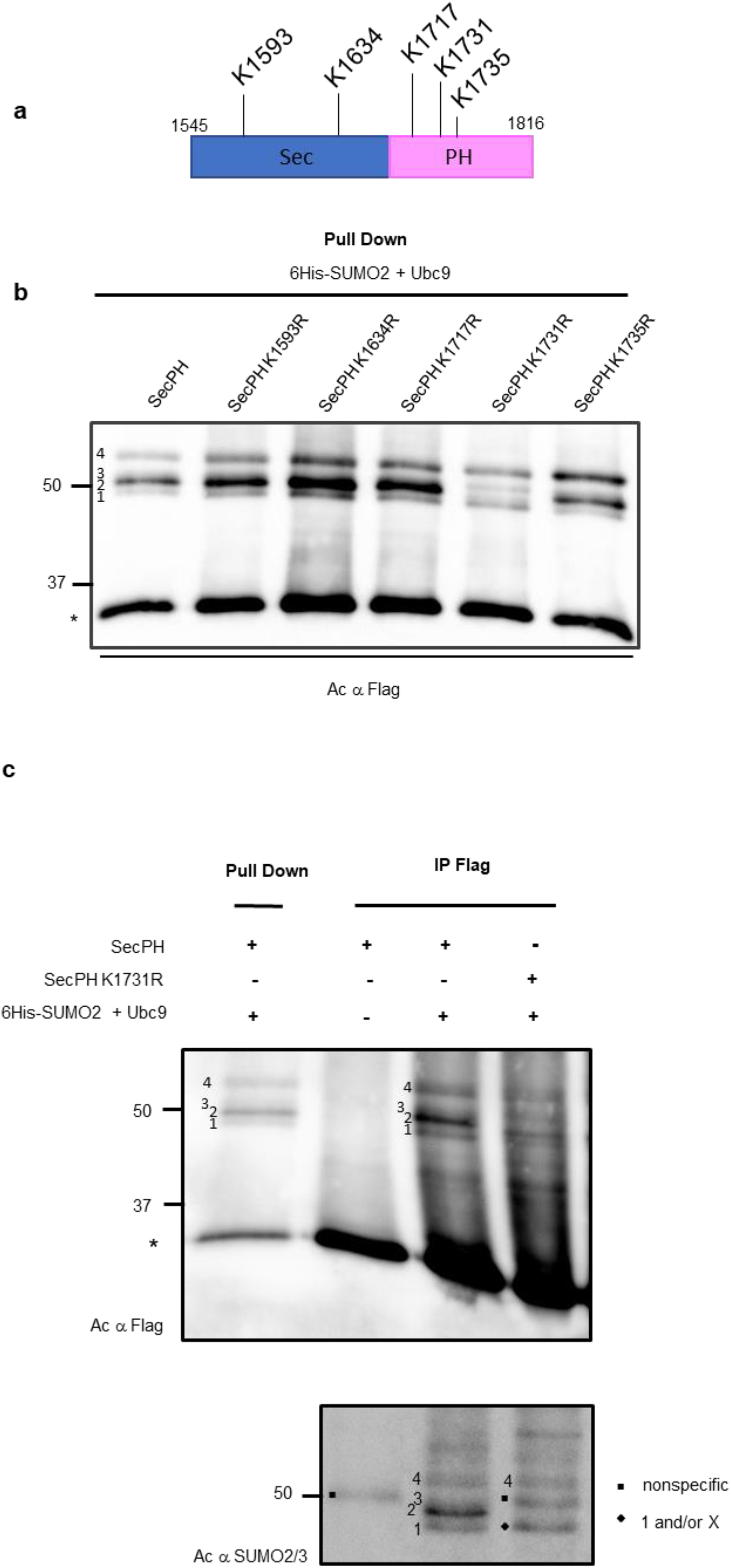
K1731 is involved in SecPH sumoylation. **a** Schematic representation of Sec and PH domains of human Nf1 (in blue and pink, respectively) and the position of the bioinformatically-predicted putative SUMOylated lysines. **b** HEK293 cells were co-transfected with pcDNA3 6His-SUMO2, pcDNA3 Ubc9, and p3XFlag SecPH or p3XFlag SecPH derivatives in which one of the five lysine was individually substituted with R (p3XFlag SecPH K1593R, p3XFlag SecPH K1634R, p3XFlag SecPH K1717R, p3XFlag SecPH K1731R, and p3XFlag SecPH K1735R). After 24 hours of growth, cells were processed and analysed as described in Fig. 2c. The four bands denoted 1 to 4 highlight the SUMOylated SecPH species. The nonspecific binding of unmodified SecPH to cobalt beads is marked with an asterisk. **c**. HEK293 cells were co-transfected with plasmids p3XFlag SecPH or p3XFlag SecPH K1731R and SUMO2 and Ubc9 overexpression vectors or empty vectors as negative control. Cell lysates were immunoprecipitated with anti-Flag antibodies and the eluates, along with the eluates from the pull-down experiments presented in (**b**), resolved by SDS-PAGE. The corresponding immunoblots revealed with anti-Flag (upper panel) and anti-SUMO2/3 antibodies (lower panel) are presented. The four SecPH SUMOylated species in the p3XFlag SecPH lane are labelled 1, 2, 3 and 4, the asterisk indicates nonspecific binding of unmodified SecPH to the cobalt beads, the diamond indicates the unresolved doublet migrating as band 1, and the square marks a nonspecific band.

To ensure that bands 1, 2, 3, and 4 corresponded to SUMOylated SecPH and not to post-translationally modified SecPH interacting with SUMO2 or SUMO2-modified proteins, we immunoprecipitated Flag-tagged SecPH and SecPH K1731R from lysates of HEK293 cells transfected with the corresponding p3XFlag SecPH vectors, pcDNA3 Ubc9, and pcDNA3 6His-SUMO2 and revealed the proteins in western blots with anti-Flag and anti-SUMO2/3 antibodies. First, the anti-Flag antibodies revealed the same pattern of bands for Flag-tagged immunoprecipitated and 6His-tagged pulled-down WT SecPH (Fig. 3c upper panel). Second, the four-band profile revealed for WT SecPH with the anti-Flag and anti-SUMO2/3 antibodies around 50 kDa was very similar (Fig. 3c lower panel), strongly suggesting that these bands correspond to SUMO-modified SecPH. Third, blotting the SecPH K1731R immunoprecipitates with the anti-Flag and anti-SUMO2/3 antibodies showed the SUMOylated band migrating as band 2 to disappear from this mutant (Fig. 3c upper and lower panel). Therefore, it is clear that bands 2 and 4 correspond to SUMOylated forms of SecPH, whereas resolution of the gels was not sufficient to separate bands 1 and x (marked with a diamond).

Immunoprecipitation of WT SecPH from cells with endogenous levels of SUMO2 and Ubc9 and long exposure of the anti-Flag immunoblot showed a band migrating at the same molecular weight as band 2, indicating that this major SUMOylation, dependent on K1731, may take place at endogenous SUMO levels (Supplementary Fig. 2a).

It has been shown that lysine residues can act as SUMO conjugation sites to compensate for each other^40^. Thus, we created a form of SecPH in which all lysines in the consensus sites were mutated to arginine (SecPH K1593,1634,1717,1731,1735R, called SecPH 5K-R) and tested it for SUMOylation. The SUMOylation profile of this mutant was similar to that of the SecPH K1634,1731R double mutant, showing stronger bands 1 and 4 and band x (Supplementary Fig. 2b). This result suggests that none of the five mutated lysines play a role in the SUMOylation events that gave rise to bands 1, 4, and x.

### 4. K1634 is a minor and K1731 a major site of SUMOylation in SecPH

We constructed a SecPH mutant with all its lysines (a total of 21) mutated into arginines (SecPH K0) and then reintroduced K1731 and K1634 into this mutant and studied their SUMOylation pattern to reinforce our results and unambiguously establish that K1731 and K1634 are SUMOylated and do not play an indirect role in the SUMOylation of other SecPH lysines. Surprisingly, SecPH K0 harboured a doublet of SUMOylation bands migrating at the same position as bands 1 and x, indicating that two lysines of the 3XFlag-tag must have been SUMOylated (Fig. 4a). As the JASSA software predicted the two lysines of the last Flag tag to be in putative SUMOylation sites (a strong inverted consensus for K4 and a weak consensus direct for K3) (Fig. 4b), we replaced them with arginines in the construct encoding WT SecPH (FlagK3K4R SecPH) and SecPH K0 (FlagK3K4R SecPH K0). Consistent with our hypothesis, band 1 and x disappeared from FlagK3K4R SecPH, whereas FlagK3K4R SecPH K0 showed no SUMOylation bands (Fig. 4c). Therefore, bands 1 and x rely on K3 and K4 of the Flag-tag. Reintroduction of K1731 into FlagK3K4R SecPH K0 mutant resulted in a new SUMOylation band migrating as band 2 (Fig. 4c) and reintroduction of both K1731 and K1634 resulted in two SUMOylation bands migrating as bands 2 and 3 (Fig. 4c) thus proving that bands 2 and 3 correspond to K1731 and K1634 SUMOylation, respectively, and that these two sites are the only ones that are detectable within the SecPH sequence under these conditions. Indeed, reintroduction of K1735, a lysine not involved in the SecPH SUMOylation pattern, into K3K4R SecPH K0 did not lead to SUMOylation (Supplementary Fig. 3).

**Fig. 4.**
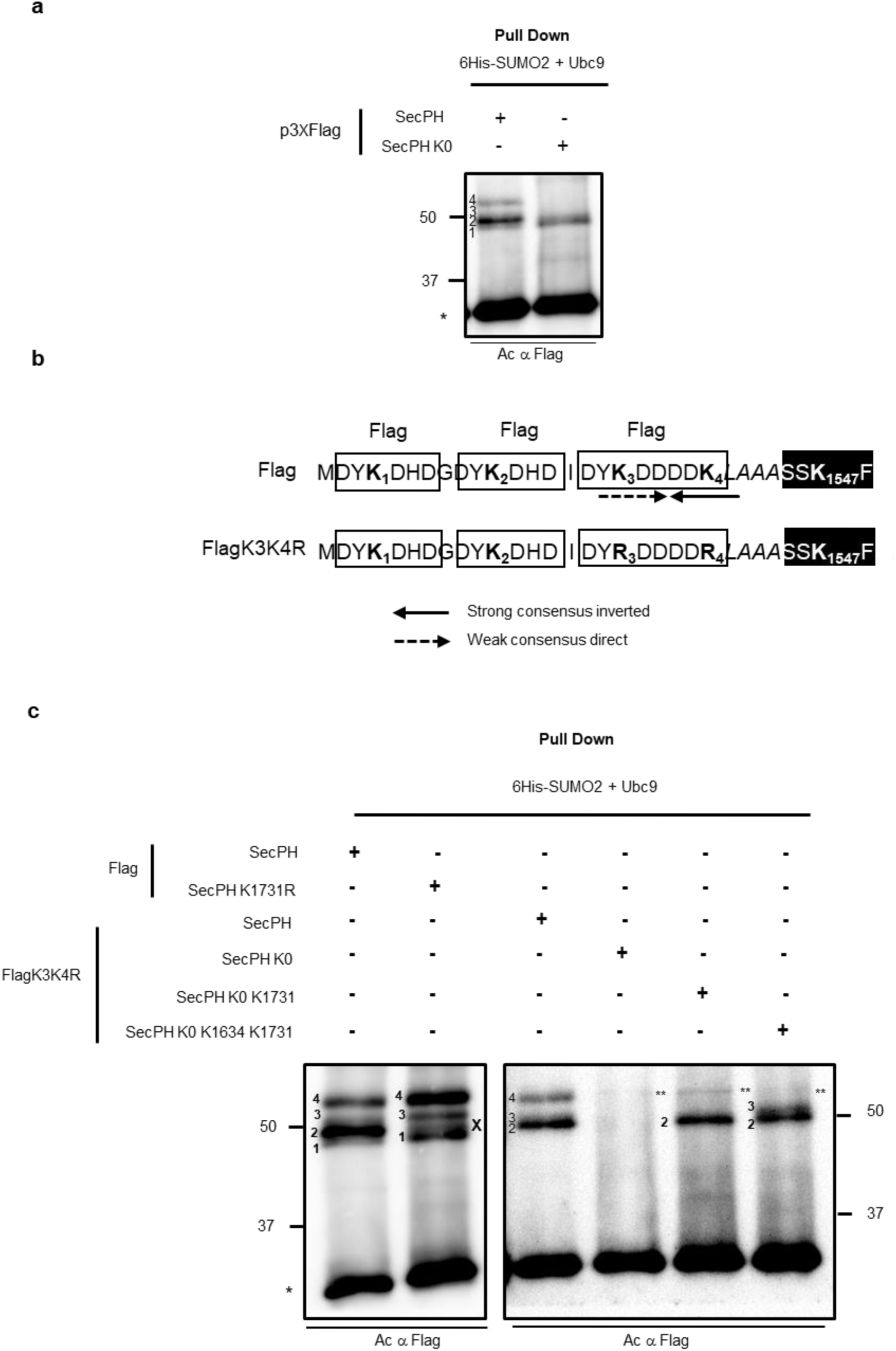
K1634 and K1731 constitute minor and major SUMO-conjugation sites in SecPH and the 3XFlag-tag accounts for another SUMO-conjugation site. **a** HEK293 cells were co-transfected with pcDNA3 6His-SUMO2, pcDNA3 Ubc9, and either p3XFlag SecPH or p3XFlag SecPH K0. Cell lysates were subjected to pull-down assays and analysed as in Fig. 2c. The asterisk (*) indicates nonspecific binding of unmodified SecPH to cobalt beads. **b** Partial amino acid sequence of p3XFlag SecPH and p3XFlagK3K4R SecPH. The three Flag sequences are boxed with the four lysines numbered sequentially, the linker is in italics, and the N-terminus of SecPH fused in frame is represented by a black box. The SUMOylation consensus sites around K3 (weak and direct) and K4 (strong and inverted) are highlighted by a direct dashed arrow and an indirect solid arrow, respectively. **c** HEK293 cells were co-transfected with pcDNA3 6His-SUMO2, pcDNA3 Ubc9, and SecPH derivatives constructed in a p3XFlag plasmid harbouring a KR substitution at the two bioinformatically-predicted SUMOylation sites K3 and K4: p3XFlagK3K4R SecPH, p3XFlagK3K4R SecPH K0, p3XFlagK3K4R SecPH K0 K1731, and p3XFlagK3K4R SecPH K0 K1634 K1731. Cell lysates were subjected to pull-down assays and analysed as in Fig. 2c. The left panel shows a control immunoblot of a pull-down assay performed with WT SecPH and SecPH K1731R cloned into a non-mutated p3XFlag plasmid, as in Fig. 2c; the right panel shows an immunoblot of the pull-down assays performed with the p3XFlagK3K4R SecPH plasmid and derivatives. Bands 1, 2, 3, 4, and x are indicated for each panel. Of note, the double asterisk (**) indicates a nonspecific band migrating slightly above band 4. The asterisk (*) indicates nonspecific binding of unmodified SecPH to the cobalt beads.

We then examined the 3D structure of SecPH^21^ and Nf1 dimer^15^ to visualize the location of K1634 and K1731. Both K1634 and K1731 are located on the surface of the SecPH 3D structure but on opposite sides (Fig. 5). In the context of Nf1 dimer, K1731 is located at the surface and accessible whether the molecule is in open or closed conformation. In contrast, K1634 is only accessible when the molecule is in open conformation (Supplementary Fig. 4).

**Fig. 5.**
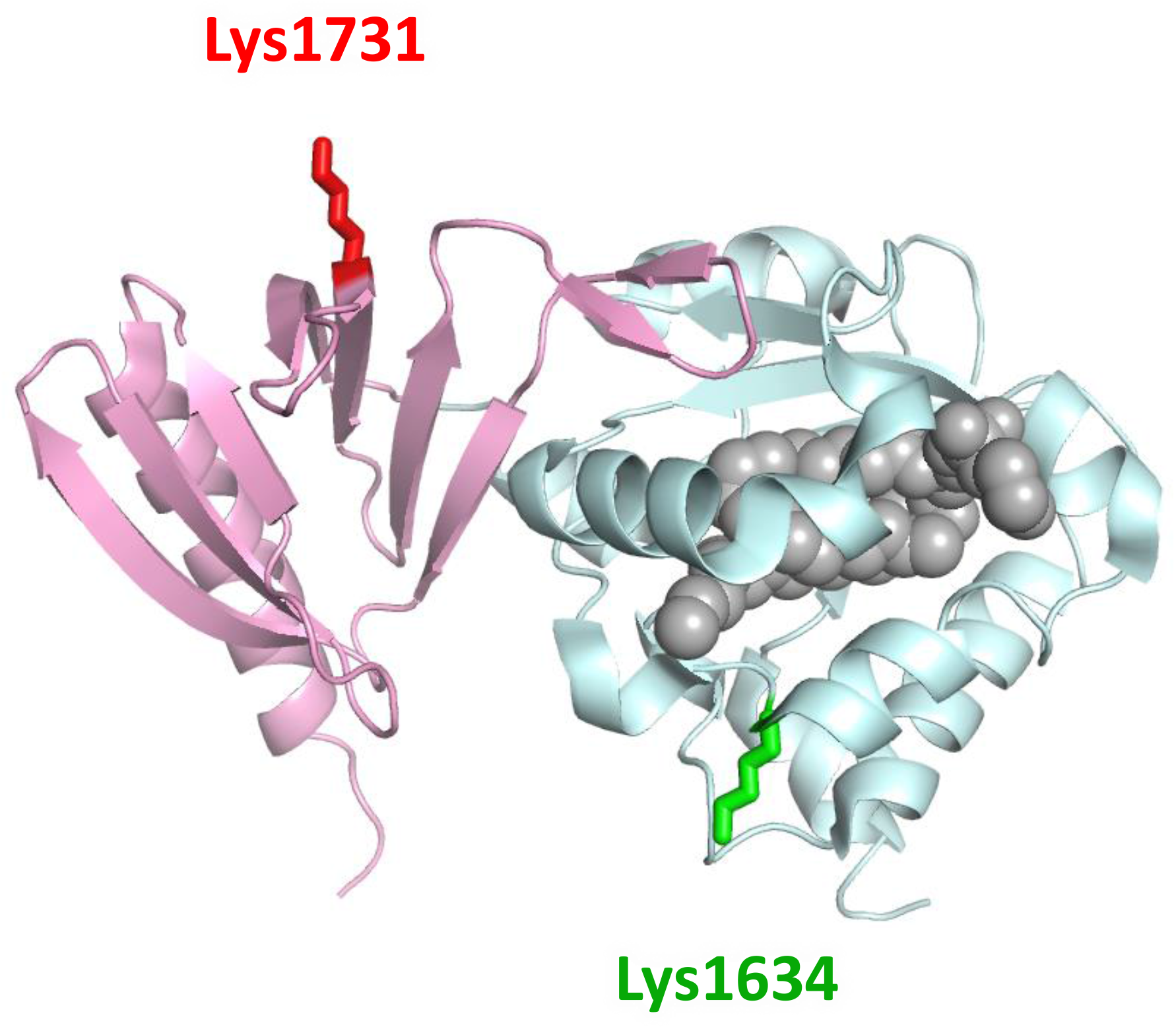
Positions of K1634 and K1731 SUMO-conjugated lysines in the 3D structure of SecPH. Ribbon representation of the neurofibromin-SecPH 3D structure complexed with phosphatidylethanolamine (with sphere representation) (PDB: 2E2X), with Sec shown in light blue, PH in pink, and phosphatidylethanolamine in grey. The K1634 and 1731 side chains are represented by sticks and labelled in green and red, respectively.

As K1731 appeared to be a strong SUMOylation site and always accessible whatever Nf1 conformation, we focused on this major SUMOylation site for the rest of the study.

### 5. K1731 SUMOylation modulates the Ras-GAP activity of the Nf1 GRD domain adjacent to SecPH

Allosteric regulation of the GAP activity of the GRD domain is possible and was demonstrated many years ago upon phosphorylation of the CSRD domain of Nf1, located just upstream of GRD^24^. By analogy, we wanted to know whether the major SecPH SUMOylation on K1731 affects GRD Ras-GAP activity. We therefore created two constructs encoding GRD-SecPH and GRD-SecPH K1731R and transfected them into HeLa cells knocked down for *NF1* (HeLa NF1^-/-^). Subsequently, we evaluated Ras-pathway activation, which is inversely proportional to Ras-GAP activity, by measuring P-ERK levels and calculating the P-ERK/ERK ratio. Ras proteins have been shown to be activated by SUMOylation^41^. Thus, we did not stimulate SUMOylation by transfecting cells with Ubc9 and SUMO2 in these experiments to avoid changes due to Ras SUMOylation.

The expression of the two constructs was similar as checked by anti-Flag western blotting (Fig. 6a) and comparison of their SUMOylation profiles showed the loss of a mono-SUMOylation band in the GRD-SecPH K1731R mutant (Fig. 6b). Quantification of the P-ERK/ERK ratio showed the GRD-SecPH construct to more efficiently compensate the loss of Nf1 and negatively regulate the RAS pathway to a higher extent (+15%) than the GRD-SecPH K1731R construct (Fig. 6c). Although this difference was small, it was significant. Only a small fraction of GRD-SecPH was SUMOylated in cells when Ubc9 and SUMO2 were not overproduced. This implies that the SUMOylated version of the protein may be considerably more active than the mutant form in stimulating Ras-GAP. Therefore, our data show the K1731R mutation, which impairs the major SecPH SUMOylation site, negatively affects the Ras-GAP activity of the Nf1 GRD domain. Given the close similarity of the K and R residues, this strongly suggests that K1731 SUMOylation stimulates the Ras-GAP activity of the Nf1 GRD domain.

**Fig. 6.**
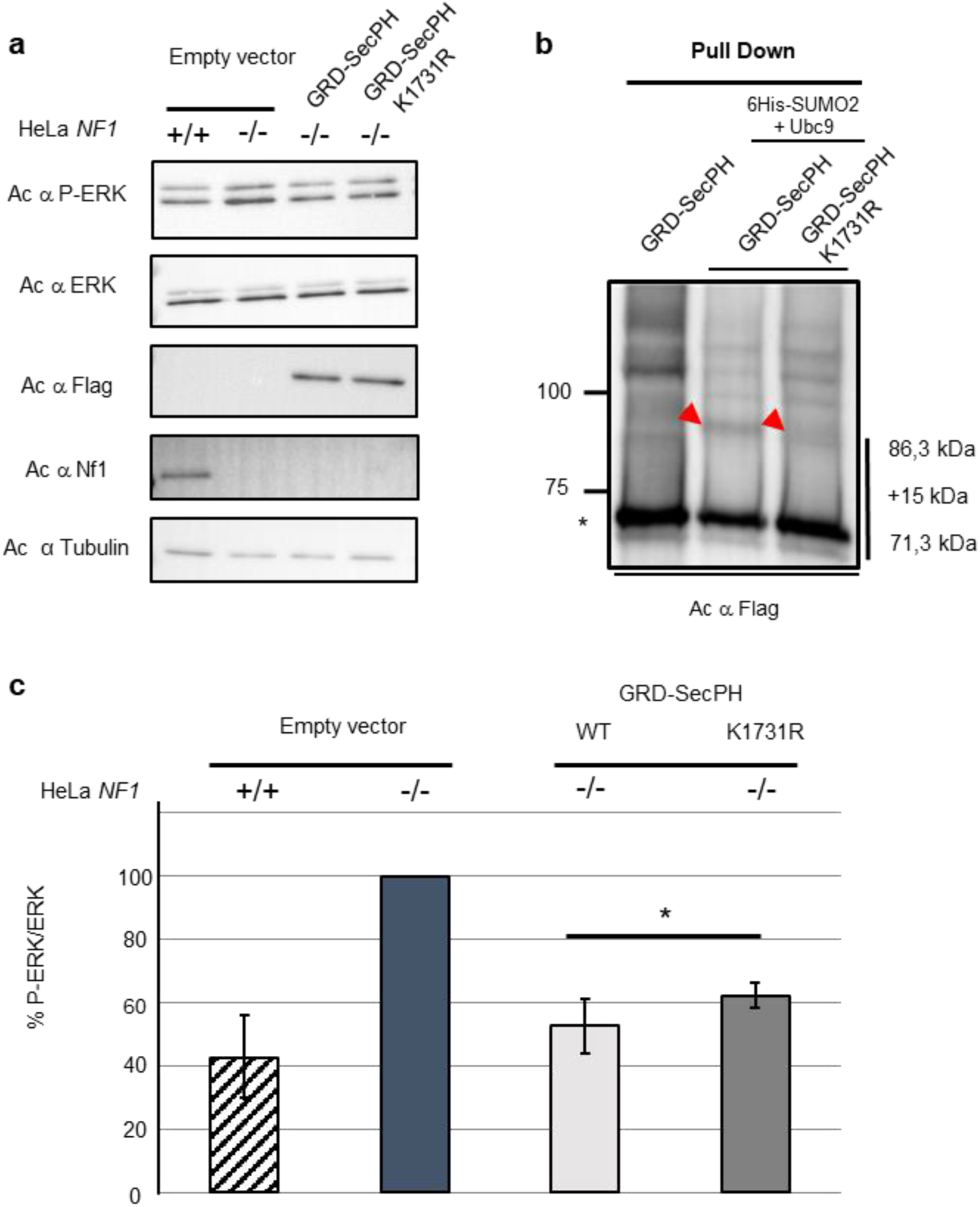
The K1731R mutation negatively affects the Ras-GAP activity of a GRD-SecPH fragment of Nf1. **a** HeLa NF1^-/-^ cells were transfected with the p3XFlag, p3XFlag GRD-SecPH, or p3XFlag GRD-SecPH K1731R plasmids and WT HeLa cells were transfected with p3XFlag. Cell lysates were analysed by western blotting using anti P-ERK, anti ERK, anti-Flag, and anti-Nf1 antibodies and anti-alpha tubulin to verify equal protein loading. Representative immunoblots are shown. **b** HEK293 cells were transfected with p3XFlag GRD-SecPH or co-transfected with pcDNA3 6His-SUMO2, pcDNA3 Ubc9, and p3XFlag GRD-SecPH or p3XFlag GRD-SecPH K1731R. Cell lysates were prepared and subjected to pull-down analysis as described in Fig. 2c. Immunoblots revealed using anti-Flag are shown. The red arrow indicates the mono-SUMOylation band that is lost in the GRD-SecPH K1731R mutant. The asterisk indicates nonspecific binding of unmodified SecPH to cobalt beads. **c** The P-ERK and ERK ratios were calculated after quantification of the immunoblots by densitometry using GeneTools from Syngene and the graphs plotted considering the P-ERK/ERK ratio of the HeLa NF1^-/-^ p3XFlag condition as 100%. Each value represents the mean ± SD of four independent experiments. Statistical analyses were performed with graphpad (* p<0,05).

### 6. The SecPH SUMOylation profile is affected in several pathogenic missense mutations

As K1731 SUMOylation appears to be important for Nf1 function, we tested whether the SUMOylation profile of NF1 patients with missense or short deletion mutations was affected. Thus, we introduced c.4733C>T (p.Ser1578Phe)^42^, c.4750A>G (p.Ile1584Val) ^43^, c.4805T>G (p.Leu1602Arg) ^44^, c.4868A>G (p.Asp1623Gly)^45^, c.4981T>C (p.Cys1661Arg)^46^, c.5205+5G>T (p.Phe1719_Val1736del) ^45^, c.5290G>T (p.Ala1764Ser)^47^, c.5360C>T (p.Thr1787Met)^48^, and c.5425C>T (p.Arg1809Cys)^49^ into our p3XFlag SecPH plasmid and studied their SUMOylation profiles. As a control, we introduced a non-pathogenic mutation, c.4972A>G (p.Ile1658Val), present in 3% of the Hispanic population^50^, into p3XFlag SecPH. Mutants S1578F, L1602R, D1623G, C1661R, and Δ1719-36 harboured a similar SUMOylation profile that was different from that of WT and the K1731R mutant (Fig. 7). This profile was characterized by a less intense band for SecPH and the presence of multiple high molecular weight bands, suggesting strong polySUMOylation. Alternatively, this pattern could correspond to mono/oligo-SUMOylated SecPH that was additionally modified by polyubiquitylation. Interestingly, the Δ1719-36 mutant, in which K1731 is deleted, exhibited exactly the same profile as the other affected patient mutants, indicating that K1731 may not be SUMOylated in any of these mutants. We tested this hypothesis by introducing the K1731R mutation into these four patient mutants (S1578F, L1602R, D1623G, and C1661R). Their SUMOylation profile proved that K1731 is not SUMOylated in these mutant forms of SecPH (Supplementary Fig. 5a). Similarly, K1634R (SUMOylated in WT SecPH) and K1717R mutations (non-SUMOylated in WT SecPH) were introduced into the D1623G mutant to assess the role of these two lysines in the SUMOylation process. The K1634R mutation did not change the D1623G mutant SUMOylation pattern. On the contrary, K1717R mutation modified it, leading to the clear loss of one mono-SUMOylation band and the enhancement of others as a probable compensation mechanism (Supplementary Fig. 5b). Overall, K1731 and K1634 do not appear to be SUMOylated in these mutants.

**Fig. 7.**
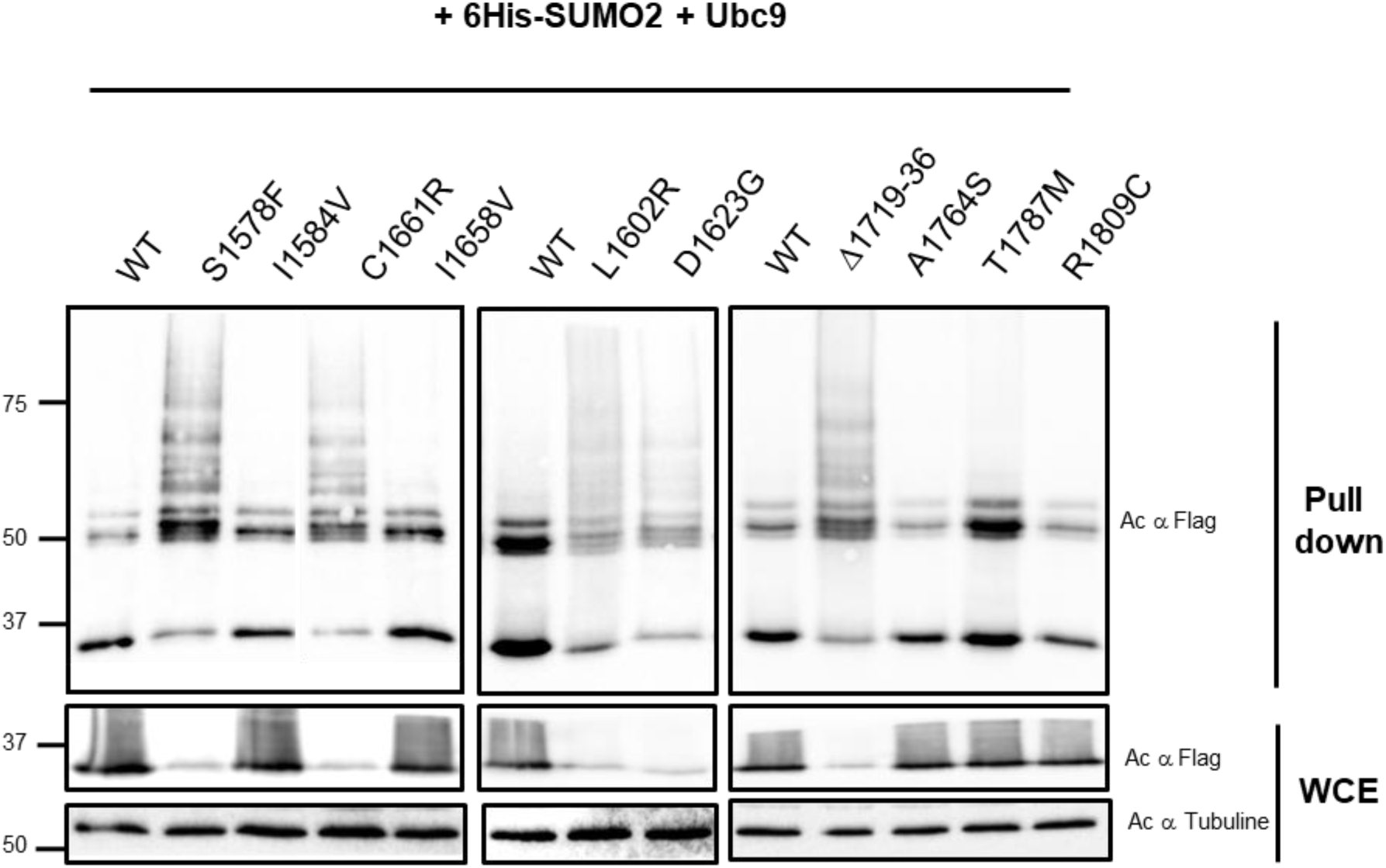
Patient missense mutations affect the SUMOylation profile and basal level of SecPH. HEK293 cells were co-transfected with pcDNA3 6His-SUMO2, pcDNA3 Ubc9, and plasmids encoding p3XFlag SecPH or p3XFlag SecPH derivatives harbouring the following patient missense mutations: S1578F, I1584V, L1602R, D1623G, I1658V, C1661R, Δ1719-36, A1764S, T1787M, and R1809C. Cell lysates were prepared and subjected to pull-down analysis as described in Fig. 2c. Immunoblots revealed with anti-Flag are shown. The different panels show different times of exposure. Cell lysates (WCE) resuspended in Laemmli buffer were analysed on immunoblots probed with anti-Flag and anti-α-tubulin antibodies as a loading control. The asterisk indicates nonspecific binding of unmodified SecPH to the cobalt beads.

When comparing mutations, we noted that mutations with an altered SUMOylation profile included a large deletion (Δ1719-36) and missense mutations of buried residues into very different, typically larger, amino-acids (S1578F, C1661R, L1602R, D1623G), all of which may be expected to result in folding defects. This hypothesis is consistent with the fact that all affected mutants showed a much lower basal level of SecPH in whole cell extracts as compared to WT SecPH (Fig. 7). By contrast, mutations that did not affect the SUMOylation pattern are either largely exposed on the surface (A1764S, T1787M, R1809C) or buried but then substituted with a similar hydrophobic residue (I1584V and I1658V). Such mutations are unlikely to disturb the folding of the domain (indeed, the 3D structure of I1584V appears to be very similar to that of WT SecPH^38^) and may instead cause disease by affecting interactions with other proteins/molecules. In conclusion, it is likely that the altered SUMOylation pattern of SecPH correlates with impaired folding.

### 7. K1731 SUMOylation does not require a classical linear consensus sequence but does require non-canonical structural elements

K1731 is part of a predicted consensus site, DTKV, depicted by JASSA^35^ to be a strong inverted consensus site fitting the pattern E/DxKΨ^37^. Such a linear motif would be expected to be more or equally accessible if the ternary structure of the protein is disturbed by mutation. However, contrary to this notion, K1731 appeared as not being SUMOylated in the five patient missense mutants, suggesting that the conformation of SecPH may be important for the SUMOylation of this lysine.

We constructed three different SecPH mutants that target the consensus sequence (Fig. 8a) to obtain further insights into the elements required for K1731 SUMOylation. The first mutant, D1729N, with the loss of the charge of this acidic residue, did not prevent K1731 SUMOylation (Fig. 8a, b). We then constructed D1729N S1733A. S1733, a possible phosphorylation site that may favour Ubc9 recruitment^51^, was abolished in this mutant. Again K1731 was still SUMOylated (Fig. 8a, b). Finally, we made a triple mutant D1729N V1732G S1733A, with a completely abolished consensus site, but K1731 was still SUMOylated in this mutant (Fig. 8a, b). Our data therefore show that K1731 is SUMO-modified independently of the predicted consensus site surrounding it in the primary structure.

**Fig. 8.**
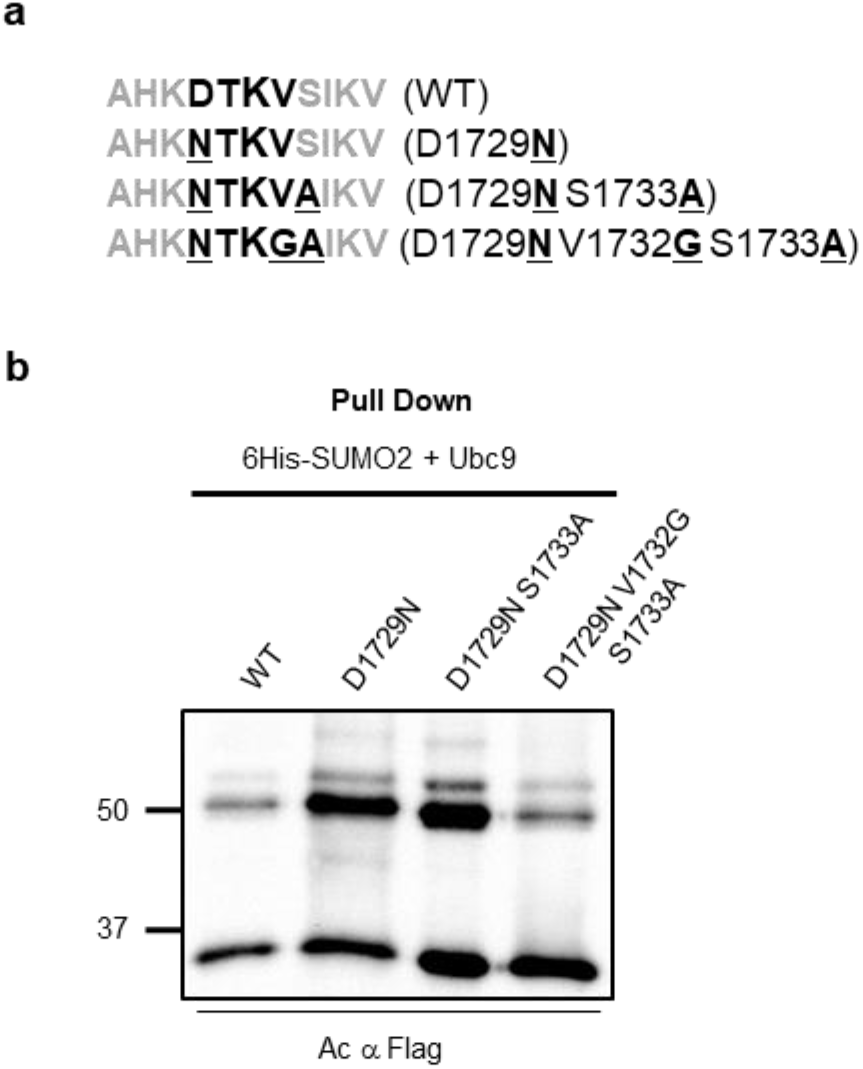
SecPH SUMOylation at K1731 is independent of the surrounding consensus site. **a** The sequence of the “predicted” inverted SUMOylation consensus site is written in black and bold and the amino acids surrounding it in grey; the major acceptor site K1731 is shown by the increased font size. The conserved amino-acid residue D1729 was replaced by N, the potentially phosphorylated S1733 mutated to A was then introduced into the D1729N variant, and, finally, a triple mutant that completely abolished the consensus site was created by changing the conserved hydrophobic residue V1732 to G in D1729N S1733A. **b** HEK293 cells were co-transfected with pcDNA3 6His-SUMO2, pcDNA3 Ubc9, and plasmids encoding p3XFlag SecPH or p3XFlag SecPH derivatives harbouring the indicated mutations. Cell lysates were prepared and subjected to pull-down analysis as described in Fig. 2c. Immunoblots revealed with anti-Flag are shown. The asterisk indicates nonspecific binding of unmodified SecPH to the cobalt beads.

JASSA software predicts, with a high score, a SUMO-interaction motif (SIM) in SecPH between residues 1621 and 1625 that is consistent with the type b consensus sequence ([P/I/L/V/M/F/Y]-[I/L/V/M]-[D]-[L]-[T])^52, 53^. Therefore, an alternative SUMOylation process could be that K1731 is SUMOylated in a SIM-dependent manner^54^. We tested this possibility by modelling (PDB: 2E2X)^21^ the interaction of the predicted SIM with SUMO from the structure of the complex between SUMO3 and the SIM of CAF1 (PDB: 2RPQ)^55^. Although SIM always interacts as a beta-strand that extends to the central beta-sheet of SUMO, the SIM of SecPH appears to be located in a middle beta-strand of a beta-sheet, and thus neighbouring beta-strands on either side may clash with SUMO (Supplementary Fig. 6). This modelling therefore strongly suggests that the predicted SIM appears to be non-functional.

We sought to obtain further insights into the structural elements required for K1731 SUMOylation. Thus, we modelled the recognition of SecPH by Ubc9 based on a structure of RanGAP1 associated with SUMO, Ubc9, and an E3 enzyme (PDB: 5D2M)^56^. Strikingly, three residues on the surface of SecPH (PDB: 2E2X)^21^, L1723, K1731, and E1747, fit into a linear consensus SUMOylation site on RanGAP1 (LK_524_xE) (Fig. 9b), suggesting that they may constitute a 3D SUMOylation consensus site, allowing an interaction with residues A131, K74, K76, and C93 of Ubc9, with adequate positioning of SecPH K1731 for SUMOylation. We then constructed simple and double L1723A E1747A mutants that would abolish this configuration. In all cases K1731 was still SUMOylated (Fig. 9c). This assumption was thus refuted.

**Fig. 9.**
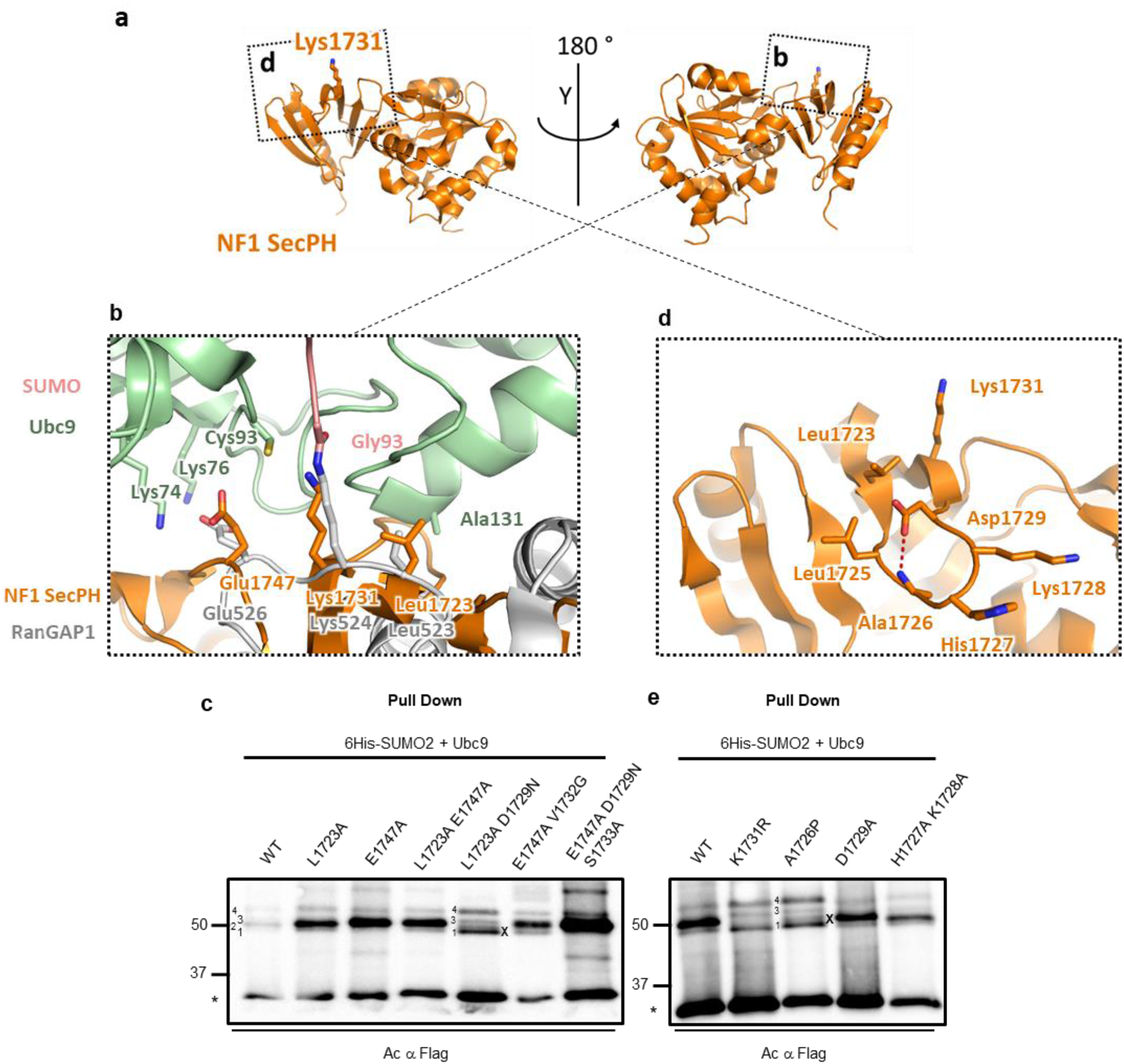
K1731 SUMOylation is independent of a putative structural consensus site in SecPH but depends on a short loop located between residues 1723-1729. **a** NF1 SecPH (PDB: 2E2X) is shown in the same orientation as in Fig. 5 and after the indicated rotation. Fragments zoomed in in (**b**) and (**d**) are marked with dashed squares. The side chain of the main SUMOylation site, Lys1731, is shown as sticks. **b** SecPH recognition by Ubc9 for K1731 SUMOylation was modelled based on a structure of RanGAP1 with SUMO, Ubc9, and an E3 enzyme (PDB: 5D2M). Close-up view of a putative structural consensus SUMOylation site surrounding K1731 fitted with the RanGAP1 consecutive consensus site. A ribbon representation is shown with RanGAP1, SUMO, Ubc9, and Nf1 SecPH coloured in grey, pink, green, and orange, respectively. Residues involved in the RanGAP1/Ubc9 interaction are labelled. Residues of the putative SecPH structural consensus site are also labelled. **c** Substitutions of amino acids belonging to the putative structural consensus site were introduced into p3XFlag SecPH and the resulting plasmids transfected into HEK293 cells, along with pcDNA3 6His-SUMO2 and pcDNA3 Ubc9. Cell lysates were subjected to pull-down analysis as described in Fig. 2c. The immunoblots revealed with anti-Flag are shown. **d** Close-up view of a short loop, located between residues L1723 and D1729. The dashed line indicates a hydrogen bond between the side chain of D1729 and the main chain of A1726. **e** Substitutions of different amino acid of the L1723-D1729 loop were introduced into p3XFlag SecPH and processed as in (**c**). The asterisk in (**c**) and (**e**) indicates nonspecific binding of unmodified SecPH to the cobalt beads.

We then hypothesized that predicted linear and postulated 3D consensus sites, or part of them, may both trigger K1731 SUMOylation. Thus, we combined mutations of their acidic or hydrophobic residues. When the E1747A mutation was combined with V1732G or D1729N S1733A, K1731 was still SUMO-modified (Fig. 9c). By contrast, combination of the L1723A mutation with D1729N resulted in the complete disappearance of the K1731 SUMOylation band and the appearance of band x (Fig. 9c). If the two apparent consensus motifs were individually functioning and compensating each other, both sets of mixed mutants should have had the same detrimental effect on K1731 SUMOylation. Therefore, these results refute the compensation hypothesis. However, the combined effect of the mutations in L1723 and D1729, which are structurally but not sequentially adjacent, underlines the importance of the structural context of K1731 for its recognition by Ubc9, consistent with the loss of K1731 SUMOylation in the folding-deficient patient mutants observed above.

Since L1723 and D1729 belong to the same short loop, we thus extended our structural investigations to this loop (Fig. 9d). It is possible that its conformation is stabilised by a hydrogen bond involving the peptide bond between L1725 and A1726 and the carbonyl of D1729, which would likely be preserved in the D1729N mutant (Fig. 9d). We constructed mutants D1729A and A1726P to abolish this hydrogen bond. Introduction of a proline in a geometrically constrained loop environment in mutant A1726P likely had an additional disruptive effect. K1731 SUMOylation appeared to be affected in the A1726P but not D1729A mutant (Fig. 9e), demonstrating the dispensability of the hydrogen bond between A1726 and D1729 but, nonetheless, confirming the importance of the loop between residues 1723-1729 for the efficient positioning of Ubc9 on SecPH. We went on to test the importance of residues located at the tip of the loop, H1727 and K1728, by mutating them to A. These substitutions had no effect on K1731 SUMOylation, suggesting that it may be the conformation of the loop rather than the specific side chains that contribute to recognition by Ubc9 (Fig. 9e).

Considering the SecPH/Ubc9-SUMO model, two other more distant structural elements at the SecPH surface may play a role in its interaction with Ubc9 (Fig. 10a). First, we probed the importance of the β-protrusion, with its two positively charged residues R1748 and K1750, which may interact with a cluster of negatively charged residues on Ubc9 (E98, E99, D100) (Fig. 10b). K1731 SUMOylation appeared to be affected in both the SecPH R1748A and SecPH K1750A mutants (Fig. 10d). These results show that the β-protrusion is crucial for K1731 SUMOylation. Second, we considered the role of two short surface loops (delineated by residues D1774 to Q1777 and M1792 to E1795) on the opposite side of K1731 that contain polar residues (E1775, N1776, and Q1794), which could interact with residues of a Ubc9 α-helix (Y134, Q139) (Fig. 10a, c). The triple mutant SecPH E1775A N1776A Q1794A had a WT SUMOylation pattern (Fig. 10d). These data suggest that these two short loops are not required for K1731 SUMOylation.

**Fig. 10.**
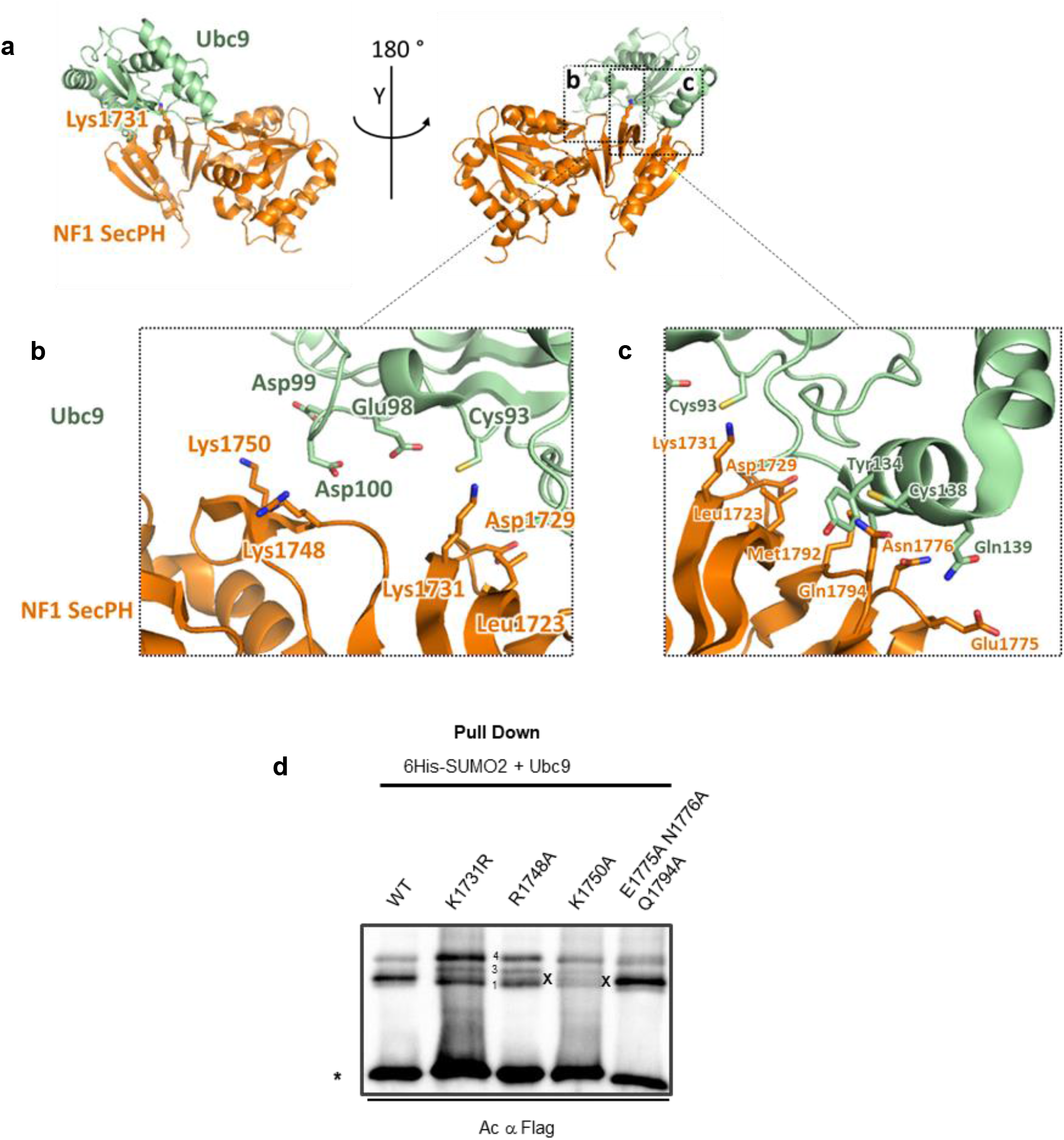
K1731 sumoylation is dependent on positively-charged residues of the β-protrusion. **a** Overall representation of the SecPH:Ubc9 interaction modelled as described in Methods. The side chain of the main SUMOylation site, Lys1731, is shown as sticks. SecPH is shown in the same orientation as in Fig. 5 and after the indicated rotation. Fragments zoomed in in (**b**) and (**c**) are marked with dashed squares. **b** and **c** Putative contacts between Ubc9 and NF1 SecPH residues of the protrusion and two short surface loops of SecPH (residues D1774 to Q1777 and M1792 to E1795). Residues possibly involved in SecPH:Ubc9 contacts are labelled and their side chains are shown. **d** Substitutions at amino acid belonging to these structural elements were introduced into p3XFlag SecPH and the resulting plasmids were transfected into HEK293 cells along with pcDNA3 6His-SUMO2 and pcDNA3 Ubc9. Cell lysates were submitted to pull-down analysis as described in Fig. 2c. The immunoblots revealed with anti-Flag are shown. The asterisk is nonspecific binding of unmodified SecPH to cobalt beads.

In summary, K1731 SUMOylation is independent of the strong linear inverted consensus sequence surrounding it. It depends on non-canonical structural elements: the conformation of a neighbouring surface loop and two positively charged residues of its β-protrusion.

## DISCUSSION

This is the first biochemical study to demonstrate that Nf1 is a SUMOylation target preferentially SUMOylated by SUMO2. This is in accordance with the fact that certain SUMO substrates are preferentially conjugated to SUMO2/3 or SUMO1, whereas others can be conjugated by either^57^. However, we cannot exclude that conjugation of Nf1 to SUMO1 is below the detection threshold of our experiments.

We showed that the Nf1 SecPH domain is SUMOylated and identified a major and a minor SUMOylation site among the five predicted consensus sites of this domain. We then demonstrated the importance of the major SecPH SUMOylation site, K1731, for Nf1 function in two ways: (i) we established the impact of this lysine on wildtype Ras-GAP activity and (ii) showed that five of nine NF1 pathogenic variants localised to SecPH that we tested show a modified SUMOylation pattern.

The physiological importance of SecPH SUMOylation at K1731 is strengthened by the existence of a c.5192A>G (p.Lys1731Arg) missense variant of *NF1* found in the gnomAD database associated with two NF1-patients for which the *in silico* predictions of deleterious effects by SIFT^58^ and PolyPhen-2 ^59^ software were inconclusive. Indeed, our results showing K1731 SUMOylation provide an unsuspected molecular explanation for the physiopathology of this mutant. Moreover, the fact that K1731 is highly conserved in most metazoan Nf1 orthologues further highlights its importance (see Supplementary Fig. 7 for a 3D structure highlighting the evolutionary conservation of Nf1 prepared using ConSurf^60^).

The localisation of K1731 within the SecPH 3D structure sheds light on how its SUMOylation could significantly affect SecPH function. SecPH is a bipartite phospholipid-binding module composed of Sec, a yeast Sec14 homologous module, that has a cavity capable of accommodating various lipids (PtdEtn, PtdGro, and glycerophospholipids)^21^. This cavity is closed by a lid-helix that is maintained in this conformation by its strong interaction with a β-protrusion of the PH domain^14^. It was hypothesized that certain events, such as ligand binding to the PH domain, could induce conformational rearrangement of the β-protrusion, possibly liberating the lid-helix of Sec and allowing its cavity to adopt an open, ligand-accepting conformation^14^. It is thus remarkable that K1731 is located very close to the base of the β-protrusion of the PH domain and is therefore at a crucial position to play a role in this rearrangement process (Supplementary Fig.8). Indeed, its SUMO-conjugation could directly affect rearrangement of the β-protrusion, either by favouring it via the creation of new interactions with the SUMO modifier or by preventing it because of steric hindrance. Therefore, SUMOylation of K1731 may regulate the open conformation of Sec and its interaction with lipids. A reverse mechanism, by which the lid conformation regulates K1731 SUMOylation, is also conceivable. Thus, direct involvement of the β-protrusion residues R1748 and K1750 in K1731 SUMOylation may offer a mechanism by which SUMOylation could act as a readout of lid conformation and, by extension, lipid binding.

Although the role of the Sec/lipid interaction remains elusive, it could target Nf1 to specific subcellular localisations to selectively regulate different Ras pathways^21^. Lipid-mediated inhibition of Nf1 Ras-GAP activity has already been reported, but this is restricted to the GRD^61^ and a similar mechanism mediated by SecPH has not been investigated. It is possible that K1731 SUMOylation has a direct allosteric effect on the GRD by affecting the Ras-GTP/GRD interaction or modifies SecPH interactions with protein partners or yet unknown properties that might influence GRD Ras-GAP activity.

Further studies are needed to clearly understand the role played by SecPH in Nf1 function to fully reveal the influence of SecPH SUMOylation.

Our results, which support a stimulatory role of K1731 SUMOylation on GRD Ras-GAP activity, do not contradict those of previous studies, which reported that SUMO2 overproduction results in Ras activation^41^. Indeed, from a physiological point of view, the regulation of Ras proteins at two levels in opposite ways could allow differential regulation of Nf1-dependent and independent Ras pathways. Other Nf1 PTMs could add further levels of complexity to the regulation of Ras pathways.

It is truly striking that five of the nine NF1 pathogenic mutants that we tested show the same SUMOylation pattern, excluding K1634 and K1731 from the SUMOylated lysines but including K1717 as a new SUMOylation site. Molecular details of these mutations versus those with a WT-like SUMOylation pattern show that changes in SUMOylation correlate with a potential folding defect and could be used in the future to discriminate between mutants that cause disease by destabilising the SecPH domain and those that involve a different mechanism. Moreover, all of the affected mutants showed a low basal level of expression, which could simply be due to their increased misfolding but could also reflect a causal link. Indeed, such an altered SUMOylation pattern might promote proteasomal degradation, possibly via the action of SUMO-targeted E3 ubiquitin ligases (STUbLs), such as RNF4 or RNF111^62^. Of note, although several E3 ubiquitin ligases have been suggested to mediate Nf1 ubiquitylation and degradation in various physiological contexts^12^, the patient mutants likely use a different pathway tailored to folding-deficient proteins and possibly involving STUbLs, which have been implicated in handling misfolded and aggregation-prone proteins^63^. The instability of SecPH missense mutants could extend to the entire Nf1 protein and thus account for their loss of function.

The absence of K1731 SUMOylation in five out of the nine pathogenic missense mutants tested prompted us to probe the role of the K1731 consensus sequence, which proved to be unnecessary for K1731 SUMOylation. Instead, we provide strong evidence that K1731 SUMOylation depends on the conformation of a neighbouring surface loop and two positively charged residues (R1748 and K1750) of the SecPH β-protrusion. Based on the 3D structure of the SecPHΔK1750 mutant, K1750 and its neighbour, R1748, have been hypothesized to be involved in the interaction of SecPH with potential ligands^38^. In light of our results, Ubc9 could be one of these ligands, and a cluster of negative residues, including D100, which has already been implicated in the interaction of Ubc9 with RanGAP1^64, 65^, may directly interact with SecPH R1748 and/or K1750. Therefore, K1731 SUMOylation, by its specific localisation, not only plays a role in Nf1 Ras-GAP activity but can also be used as a read out of SecPH folding for stabilisation purposes.

This study therefore shows consensus site-independent SUMOylation and provides unexpected structural information about Ubc9 positioning and binding to SecPH for effective SUMO transfer. Although a full understanding of the SecPH/Ubc9 interaction and effective SUMO transfer requires resolution of the 3D structure of the complex, our data nevertheless shed light on yet unexplained mechanisms of SUMO-conjugation.

Our study highlights SUMOylation as a novel Nf1 PTM that plays an important role in its function. In the future, this PTM should be considered in light of the recent resolution of the 3D structure of the Nf1 dimer showing the existence of inactive (closed) and active (open) conformations^15^. Other post-translational modifications of Nf1, such as phosphorylation and ubiquitylation, have already been shown to play such a role. The interplay between the SUMO and ubiquitin pathways has been demonstrated in the functional modulation of several substrates modified by these two proteins^66^. Similarly, Nf1 functions are likely orchestrated by various PTMs, and the interplay between them, including SUMOylation, warrants further investigation in the future.

## METHODS

### Recombinant DNA and site-directed mutagenesis

pcDNA3 Ubch9/SV5 tagged (thereafter named pcDNA3 Ubc9) and pcDNA3-His_6_-SUMO-2 (thereafter named pcDNA3 6His-SUMO2) were kindly provided by R. Hay and have been previously described^67, 68^. pcDNA3 6His-SUMO2 amplification with Flag SUMO2 primers (all primers are listed in Supplementary Table 1) generated an amplicon which was eventually cleaved with HindIII and BamHI and inserted into the same sites of p3XFLAG-Myc-CMV™-24 Expression Vector (E9283, Sigma-Aldrich) to create p3XFlag SUMO2 encoding a 3XFlag SUMO2 fusion protein. p3XFlag SUMO1 was synthesised at GenScript. p3X Flag SecPH^19^ was used the create most of p3XFlag SecPH plasmids derivatives with one, two, or more amino acid substitutions in SecPH with Q5 PCR methodology (Q5®Site-Directed Mutagenesis Kit, New England Biolabs, E0554) or PCR site-directed mutagenesis strategy. To construct p3XFlag SecPH 5K-R, K1731R was first introduced into p3XFlag SecPH K1735R with primers SecPH K1731,1735R; then the three other KR substitutions (K1593R, K1634R, K1717R) were introduced one after the other by successive rounds of mutagenesis. p3XFlag SecPH Δ1719-36 was serendipitously obtained during construction of a p3XFlag SecPH variant. p3XFlag SecPH A1726P, p3XFlag SecPH D1623G K1717R, p3XFlag SecPH H1727A K1728A, p3XFlag SecPH E1775A N1776A Q1794A, p3XFlag SecPH K0, p3XFlagK3K4R SecPH plasmid and p3XFlagK3K4R SecPH derivatives with various amino acids substitutions in SecPH were synthesised at GenScript.

To create p3XFlag GRD-SecPH encoding a 3XFlag GRD-SecPH fusion protein (from Gly1168 to Asp1816 in Nf1 isoform1) a stop codon was inserted into plasmid p3XFlag GRD-SecPH-EX by Q5 site directed mutagenesis and primers GRD-SecPH. The 3XFlag GRD-SecPH K1731R variant was created as above.

To generate p3XFlag GRD-SecPH-EX plasmid, an amplicon made with a library from human lung (Marathon-ReadyTM, BD Biosciences) and primers GRD-SecPH-EX was cleaved with NotI and SalI, and inserted into the identical sites of p3XFlag-Myc-CMV™-24 (ibidem). The 3XFlag GRD-SecPH-EX fusion protein extends from Gly1168 to Thr1884 in Nf1 isoform1.

MDH1-Flag was purchased from GeneCopoeia (EX-C0441-M13-10)

### Cell growth and Transfection

HEK293T cells from the American Type Culture Collection (ATCC ref: CRL1573, thereafter named HEK293) were cultured under 5% CO_2_ at 37°C in low-glucose Dulbecco’s modified Eagle’s medium (DMEM) (Sigma–Aldrich, D6046) supplemented with 10% foetal bovine serum (FBS-Gibco). Transient transfection using the calcium-phosphate method was carried out with 15 or 20 µg of plasmid/100-mm dish. Further experiments were conducted 48 h after transfection.

HeLa NF1^+/+^ and HeLa NF1^-/-^ cells (BioCat) were cultured under 5% CO_2_ at 37°C in high-glucose DMEM (Sigma-Aldrich, D5796) supplemented with 10% FBS. Transient transfection was performed using Lipofectamine 2000 (Invitrogen) and 15 µg of plasmid DNA. After 24 hours of growth, cells were serum-starved (0.1% FBS) in high-glucose DMEM for 18 h to decrease to a basal level the activation of the Ras pathway. After 18 h of growth-factor deprivation, the cells were restimulated for 30 min with fresh medium supplemented with 1% FBS immediately before the preparation of cell lysates (adapted from Cichowski ^26^).

Cells were regularly checked for mycoplasma contamination using a luminescence-based kit (Lonza, MycoAlert^TM^ Mycoplasma Detection Kit).

### Immunoprecipitation for the detection of endogenous Nf1 SUMOylation

HEK293 cells were transiently transfected with p3XFlag SUMO1 or p3XFlag SUMO2 and pcDNA3 Ubc9. After 48 h of growth, cells were lysed and the lysates enriched for SUMOylated proteins by a first IP and then a second IP was performed to capture SUMOylated Nf1.

Human HEK293 cells were seeded on three 100-mm collagen coated dishes, cultured for 24 h, and co-transfected, using the calcium phosphate method, with two expression plasmids, each encoding Flag SUMO paralogs and Ubc9. After two days of growth, cells were washed three times with cold phosphate-buffered saline (PBS) supplemented with 20 mM NEM. Cells were then lysed for 10 min on ice in 500 μl lysis buffer containing 50 mM Tris-HCl (pH 7.5), 100 mM NaCl, 5 mM EDTA, 0.1% Triton-X100, 1 mM PMSF, 1 µM E-64, 20 mM NEM, and EDTA-free protease inhibitor cocktail (resuspended as recommended by the manufacturer Sigma-Aldrich, S-8830). After centrifugation at 10,000 x g for 10 min at 4 °C, the clarified lysates were incubated with equilibrated anti-Flag M2 affinity gel (ANTI-FLAG M2 Affinity Gel Sigma-Aldrich, A2220) on a rotating wheel for 3 h at 4°C. The beads were subsequently washed five times with ice-cold lysis buffer freshly supplemented with 20 mM NEM and subsequently incubated with Flag peptide (2 mg/mL, Sigma-Aldrich, F-3290) for 30 min at 4°C. The eluted SUMOylated proteins were then mixed with 10 µl undiluted anti-Nf1 antibody (Novus Biologicals, NB100-418) and incubated for 2 h at 4°C on a rotating wheel. Then, the anti-Nf1-bound proteins were captured by an additional incubation for 1 h at 4°C with Protein G beads (Pierce, 2851), followed by extensive washing in ice-cold lysis buffer freshly supplemented with 20 mM NEM. The captured proteins were eluted in Laemmli buffer by incubation at 95°C for 5 min, followed by a centrifugation step at 10,000 x g for 5 min.

### Immunoprecipitation and affinity pull-down experiments after transfection of p3XFlag SecPH or p3XFlag GRD-SecPH and their variants into HEK293 cells

HEK293 cells were co-transfected with p3XFlag SecPH or p3XFlag GRD-SecPH or either of their variants, along with pcDNA3 6His-SUMO2 and pcDNA3 Ubc9 or the corresponding empty vectors. As a negative control, HEK293 cells were co-transfected with the MDH1-Flag plasmid, along with pcDNA3 6His-SUMO2 and pcDNA3 Ubc9. After 48 h of growth, the cells were processed as follows. For Immunoprecipitation analyses, cells were lysed for 10 min with 500 μl ice-cold lysis buffer consisting of 50 mM Tris-HCl (pH 7.5), 300 mM NaCl, 5 mM EDTA, 0.1% Triton-X100, 1 mM PMSF, 50 mM NaF, 1 µM E-64, 20 mM NEM, and EDTA-free protease inhibitor cocktail (resuspended as recommended by the manufacturer, Sigma-Aldrich, S-8830) and then centrifugated at 10,000 x g for 10 min at 4°C. Flag-tagged SecPH proteins in the cell extracts were captured with an anti-Flag M2 affinity gel (ANTI-FLAG M2 Affinity Gel, Sigma-Aldrich, A2220) by incubation on a rotating wheel at 4°C for 3 h. After extensive washing in 300 mM NaCl ice-cold lysis buffer, bound proteins were eluted with Flag peptide, as above, and the eluted proteins recovered after centrifugation. For the enrichment of SUMOylated proteins by affinity pull-down, transfected HEK293 cells were recovered by low-speed centrifugation at 4°C, washed two times in 3 ml ice-cold PBS buffer, and then resuspended in 1 ml PBS. One tenth of the cell resuspension was used to prepare whole-cell extracts with 100 µL ice-cold lysis buffer consisting of 50 mM Tris-HCl (pH 7.5), 100 mM NaCl, 5 mM EDTA, 0.1% Triton-X100, 1 mM PMSF, 50 mM NaF, and EDTA-free protease inhibitor cocktail. After 10 min incubation on ice, cell debris were removed by centrifugation at 10,000 x g for 10 min at 4°C. Then, 60 µL of the extract was mixed with 20 µL 4X Laemmli buffer and samples stored at −80°C. The second part of the cell resuspension was centrifuged at low speed and cell extracts prepared under denaturing conditions by incubation in 500 µL 6M guanidium hydrochloride buffer pH 7.6 (6M guanidium hydrochloride prepared in 100 mM phosphate buffer) at 95°C for 5 min followed by extensive vortexing to remove chromatin-bound proteins. Insoluble material was removed by centrifugation at 16,000 x g for 30 min at room temperature (RT). The supernatant was then mixed with 50 µL cobalt beads (TALON® Metal Affinity beads Clontech, 635502) prewashed three times in guanidium hydrochloride buffer and the samples incubated at RT for 3 h with rotation. Pulled-down His-tagged SUMO-conjugated proteins were washed twice in guanidium hydrochloride buffer, twice in buffer containing guanidium hydrochloride buffer and imidazole buffer (20 mM Imidazole, 25 mM Tris-HCl, pH 6.8) at a 1:3 ratio, and, finally, three times in imidazole buffer. The SUMO-conjugated proteins were eluted by incubation at 95°C for 5 min in Laemmli buffer, followed by centrifugation for 10 min at 16,000 x g. Samples were subsequently analysed on 9% SDS-PAGE alongside whole-cell extracts.

### Immunoblotting

Proteins were transferred onto Immobilon-P PVDF membranes (Merck IPVH00010) using a wet-transfer system (Bio-Rad) at 100 V and 4°C for 90 min. Then, membranes were blocked in 5% non-fat milk prepared in Tris-buffered saline with 0.1% Tween-20 (TBS-T). After 1 h at RT, the buffer was discarded and the primary antibody (listed in Supplementary Table 2), diluted into fresh TBS-T + 5% non-fat milk, was added. After incubation overnight at 4°C, the membrane was washed three times with TBS-T and the rabbit (Invitrogen, 65-6120,) or mouse (Invitrogen, 61-6520) secondary antibody conjugated to horseradish peroxidase (diluted 1:50 000 in TBS-T+ 5% non-fat milk) was added to the membrane and incubated at RT for 2 h. After three washes in TBS-T (20 min each), the proteins were visualized by chemiluminescence (SuperSignal™ West Dura Extended Duration Substrate, Pierce™) using the PXi imaging system (Syngene).

### Antibodies and reagents

The primary antibodies are listed in Supplementary Table 2.

### Ras GAP activity measurement

P-ERK and ERK levels were measured in HeLa cells co-transfected and grown as described above. Cells were lysed in 100 µL ice-cold buffer for 5 min (50 mM Tris-HCl pH 7.5, 100 mM NaCl, 5 mM EDTA, 1% Triton-X100, and protease inhibitors (50 mM NaF, 10 mM sodium pyrophosphate, 1 mM Na3VO4, 20 mM p-nitrophenyl phosphate, 20 mM β-glycerophosphate, 10 mg/ml aprotinin, 0.05 mg/ml okadaic acid, 1 mg/ml leupeptin, and 1 mM PMSF). After centrifugation at 10,000 x g for 10 min at 4°C, the total protein concentration was determined by the Bradford method using the Bio-Rad Protein Assay Kit II (Bio-Rad, 5000 002) and serial dilutions of bovine serum albumin as a calibration standard (BSA, Sigma-Aldrich). An equal amount of protein was separated on 10% or 7% SDS-PAGE (for NF1 detection), immunoblotted, and analysed using anti-ERK, anti-pERK, anti-Nf1, anti-Flag, and anti β-tubulin antibodies, as described in the previous paragraph. The P-ERK and ERK ratios were calculated after quantification of western blots by densitometry using GeneTools from Syngene. Statistical significance was determined using one-way ANOVA (***P < 0.001, **P < 0.01, *P < 0.05). Error bars correspond to standard deviations.

### Structural models and figures and sequence conservation analysis

Structural models and figures were produced in PyMOL Molecular Graphics System, Version 2.0 (Schrödinger, LLC). For all analyses, we used the crystal structure of Nf1 SecPH bound to phosphatidylethanolamine (PDB: 2E2X)^21^. Superposition with the structure PDB: 5D2M^56^ in Fig. 9b was performed by aligning Cα and Cβ atoms of indicated Glu, Lys, and Leu residues in NF1 SecPH and RanGAP1 using the command “pair fit” in PyMOL. The model of the SecPH/Ubc9 interaction in Fig. 10 was produced from the superposition in Fig. 9b by further manually moving Ubc9 to avoid clashes and increase interaction surface. The comparison between the central β-sheet of the Sec subdomain of Nf1 and a SUMO:SIM structure (PDB: 2RPQ)^55^ was performed by aligning the indicated sequence motifs using the command “super” in PyMOL. Sequence conservation analysis in Supplementary Fig.7 was performed with the HMMER method using the ConSurf 2016 software^60^ with default settings except for focusing only on homologues with at least 65% identity in the SecPH domain, which corresponds to Nf1 from most metazoan species except for distant outliers. In this group, K1731 appears strictly conserved. For the representation of sequence conservation, ConSurf conservation scores were projected onto a SecPH crystal structure (PDB: 2E2X)^21^ via the ConSurf server.

### Figure preparation

Graphs were prepared using Microsoft Excel 2016 and further edited using Power Point, which was also used to assemble all figures.

## ACKNOWLEDGEMENTS

We thank Dr. Ronald Thomas Hay (Centre for Gene Regulation and Expression, College of Life Sciences, University of Dundee, Dundee, Scotland DD1 5EH, UK) for providing the plasmids. We are grateful to laboratory alumni for their technical support (Veronika Buršić, Manon Julien, Navid Barakzoy, Aurélie Gombault, Marie-Ludivine de Tauzia, and Khadija Aziz). We thank Alessia Zamborlini, Stéphane Martin, Eric Pasmant for stimulating discussions. The CNRS, the French Association Neurofibromatose et Recklinghausen (research grant to H.B.), the Ligue Contre le Cancer (research grant to H.B. and B.V.), and the French Agence Nationale de la Recherche (CliNeNF1, grant number: ANR-19-CE18-0016-02) provided financial support. The authors gratefully acknowledge their financial contribution to conduct these studies. M.B. was supported by a fellowship from the French Ministère de l’Enseignement Supérieur, de la Recherche et de l’Innovation.

## AUTHOR CONTRIBUTIONS

M.B. and C.M. conceived and performed most of the experiments. M.B., C.M., and I.S. performed the cloning, mutagenesis, and pull-down experiments. F.G., M.D., C.M., and M.B. performed the IP experiments. M.J.S. prepared structural models and figures and proposed structure-guided mechanistic hypotheses. F.G., M.D., and M.B. contributed to the final figures and provided supporting data. B.V., C.M., and M.S. reviewed and edited the manuscript. H.B. conceived the project and H.B. and B.V. supervised the study. H.B. wrote the original draft with the assistance of B.V, M.S., and C.M. All authors contributed to revising the manuscript.

## ADDITIONAL INFORMATION

### Supplementary information

**Supplementary Figure 1.**
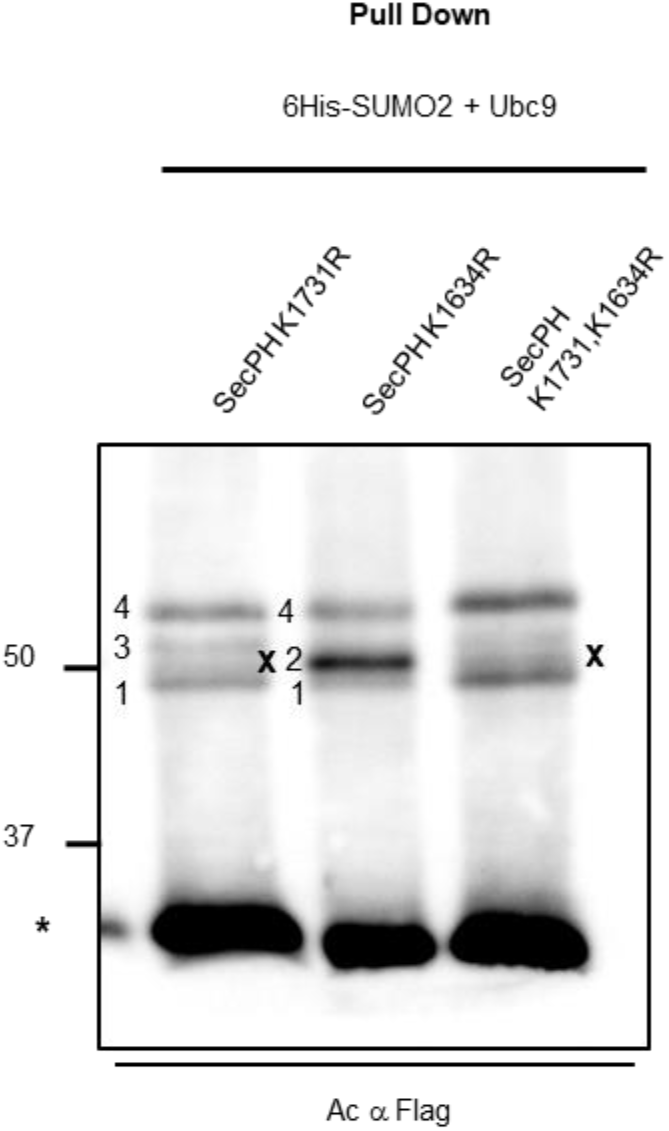
K1634 is involved in SecPH SUMOylation and a supplementary band appears in K1731R mutants (related to Fig. 3b) HEK293 cells were transiently co-transfected with plasmids encoding SecPH derivatives: p3XFlag SecPH **K1634R** or p3XFlag SecPH **K1731R or** p3XFlag SecPH **K1634,1731R,** and SUMO2 and Ubc9 overexpression vectors. Cell lysates were subjected to pull-down assays and the eluted proteins analysed as in Fig. 2c. Bands 1, 2, 3, and 4 described in Fig. 2c are shown; the new band that appears just above band 1 in p3XFlag SecPH **K1731R** and p3XFlag SecPH **K1634,1731R** is denoted « x ». The nonspecific binding of unmodified SecPH to the cobalt beads is indicated by an asterisk.

**Spplementary Figure 2.**
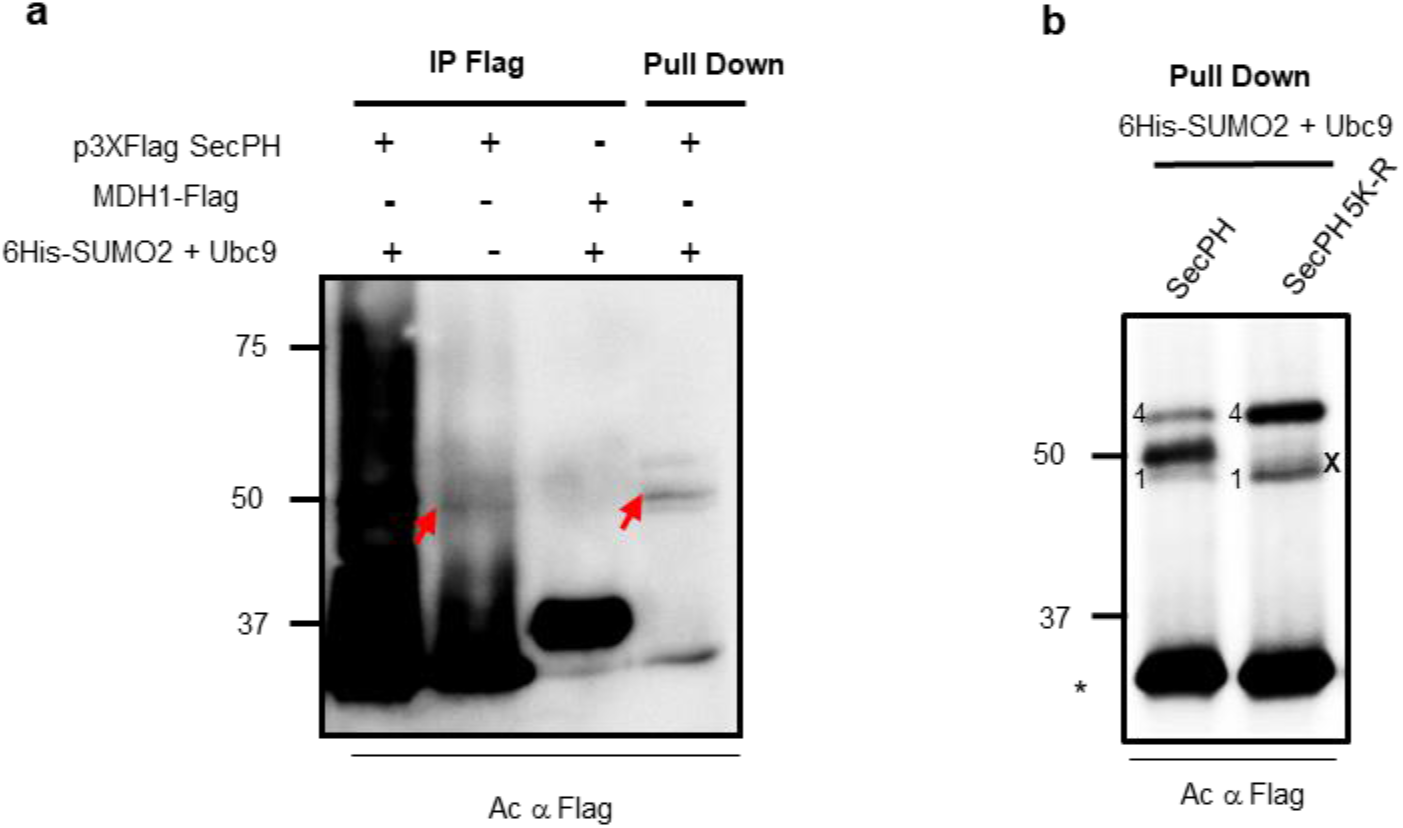
SecPH is SUMOylated at endogenous SUMO levels and other SecPH SUMOylation consensus sites are not modified by SUMO (related to Fig. 3c) **a** HEK293 cells were transfected with p3XFlag SecPH, with or without SUMO2 and Ubc9 overexpression plasmids, or with MDH1Flag as a negative control. Cell lysates were immunoprecipitated with anti-Flag antibodies and the eluates resolved by SDS-PAGE, along with eluates from the pull-down experiment presented in Fig. 3b. An over-exposure of an immunoblot revealed with anti-Flag antibodies is presented. A band migrating as band 2 in Fig. 3c is shown by a red arrow. **b** HEK293 cells were transiently co-transfected with p3XFlag plasmids encoding SecPH or the SecPH 5K-R mutant (p3XFlag SecPH K1593, 1634, 1717, 1731, 1735R) and SUMO2 and Ubc9 overexpression vectors. Preparation of lysates and pull-down assays were performed and analysed as described in Fig. 2c. The bands x, 1, 2, 3, and 4, corresponding to SecPH SUMOylated species in p3XFlag SecPH, are shown. The nonspecific binding of unmodified SecPH to the cobalt beads is indicated by an asterisk.

**Supplementary Figure 3.**
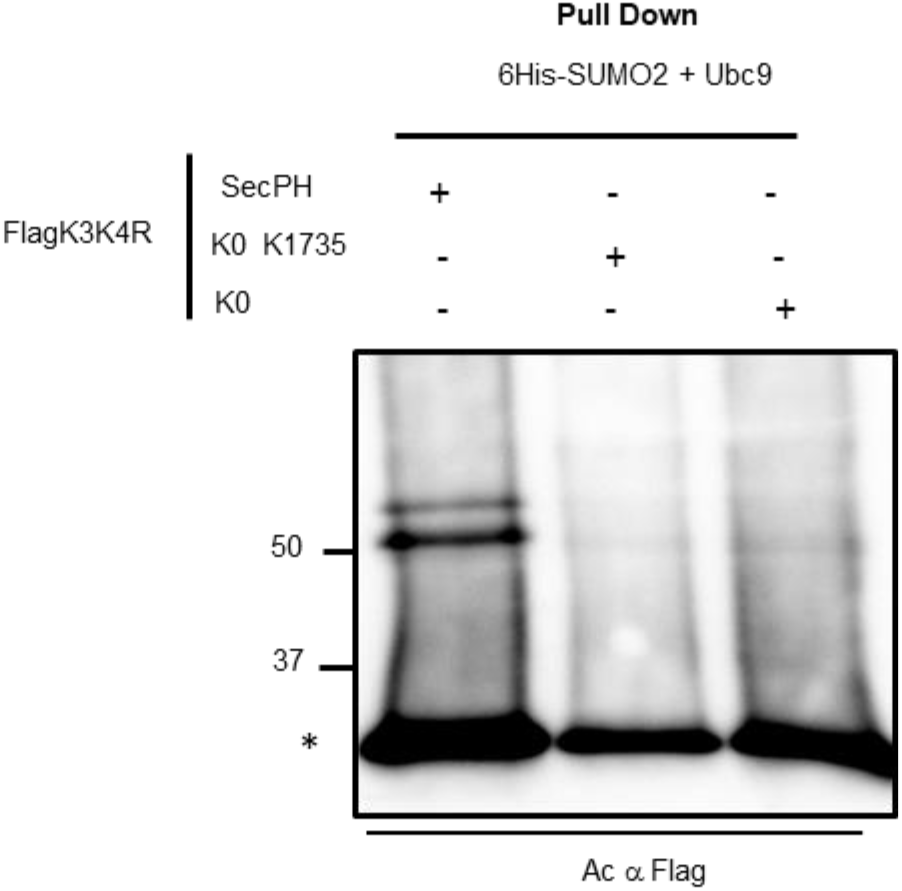
K1735 is not SUMOylated when reintroduced into a SecPH K0 mutant (related to Fig. 4c) HEK293 cells were co-transfected with pcDNA3 6His-SUMO2, pcDNA3 Ubc9, and SecPH variants (SecPH, SecPH K0, and SecPH K0 in which K1735 was reintroduced) cloned into plasmid p3XFlagK3K4R harbouring a KR substitution at the two bioinformatically-predicted SUMOylation sites K3 and K4. Cell lysates were subjected to pull-down assays and analysed as in Fig. 2c. The nonspecific binding of unmodified SecPH to the cobalt beads is indicated by an asterisk.

**Supplementary Figure 4:**
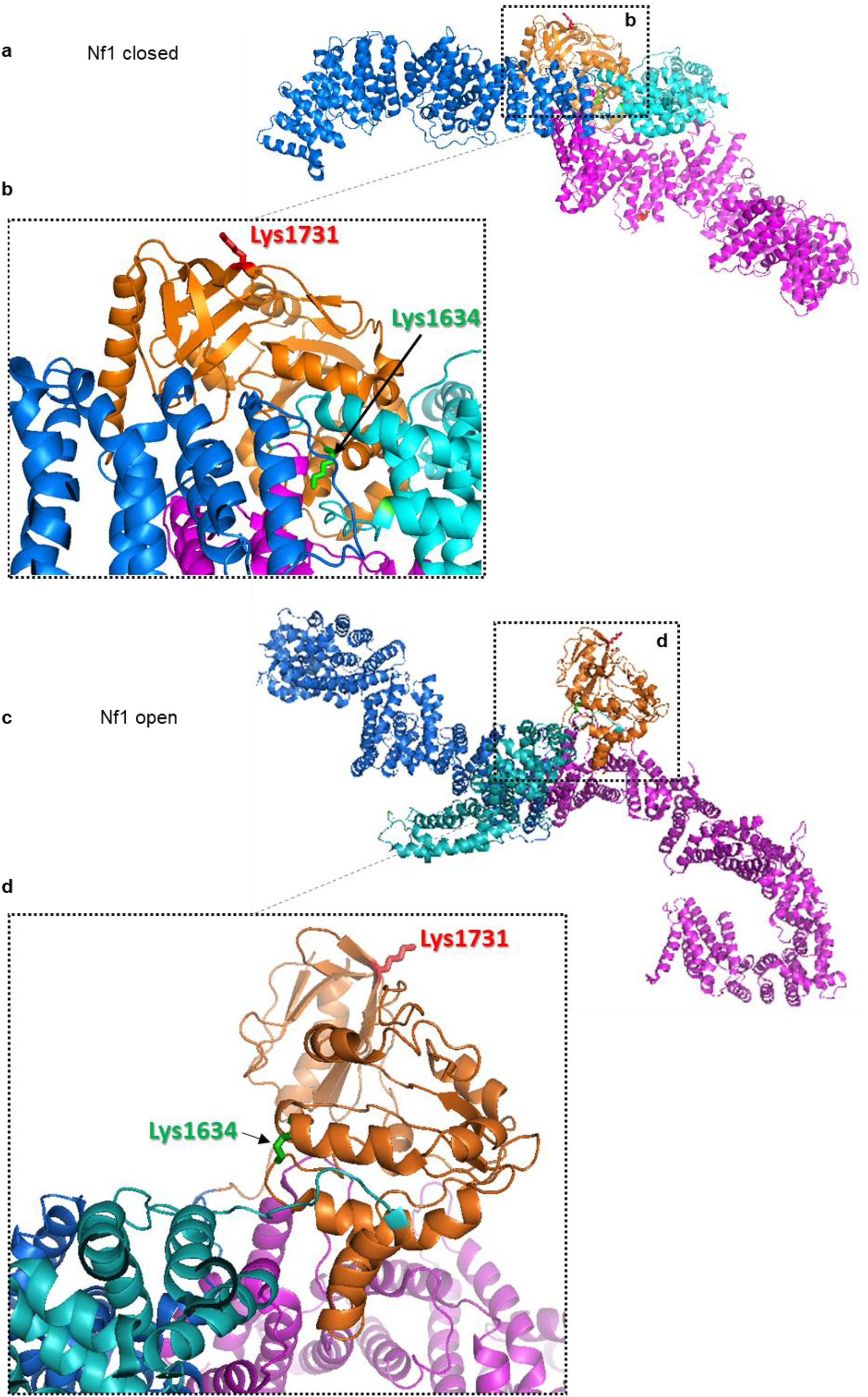
Positions of the two SUMO-conjugated lysines in Nf1 isoform2 3D-structure (related to Fig. 5) Overall ribbon representation of Nf1 isoform2 3D structure in closed (PDB: 7PGU) (**a**) and in open (PDB: 7PGT) (**c**) conformations. Nf1 domains are colored differently: N HEAT-ARM domain in magenta, GRD in Cyan, SecPH in orange and C-HEAT/ARM in marine. K1634 and K1731 side chains are represented as sticks and labelled in green and red, respectively. Residue numbering from isoform1 was used for consistency with other figures, but real residue numbers in isoform2, which contains a 21-amino-acid insertion corresponding to exon 23a, would be K1655 and K1752. Fragments zoomed in (**b**) and (**d**) are marked with dashed squares.

**Supplementary Figure 5.**
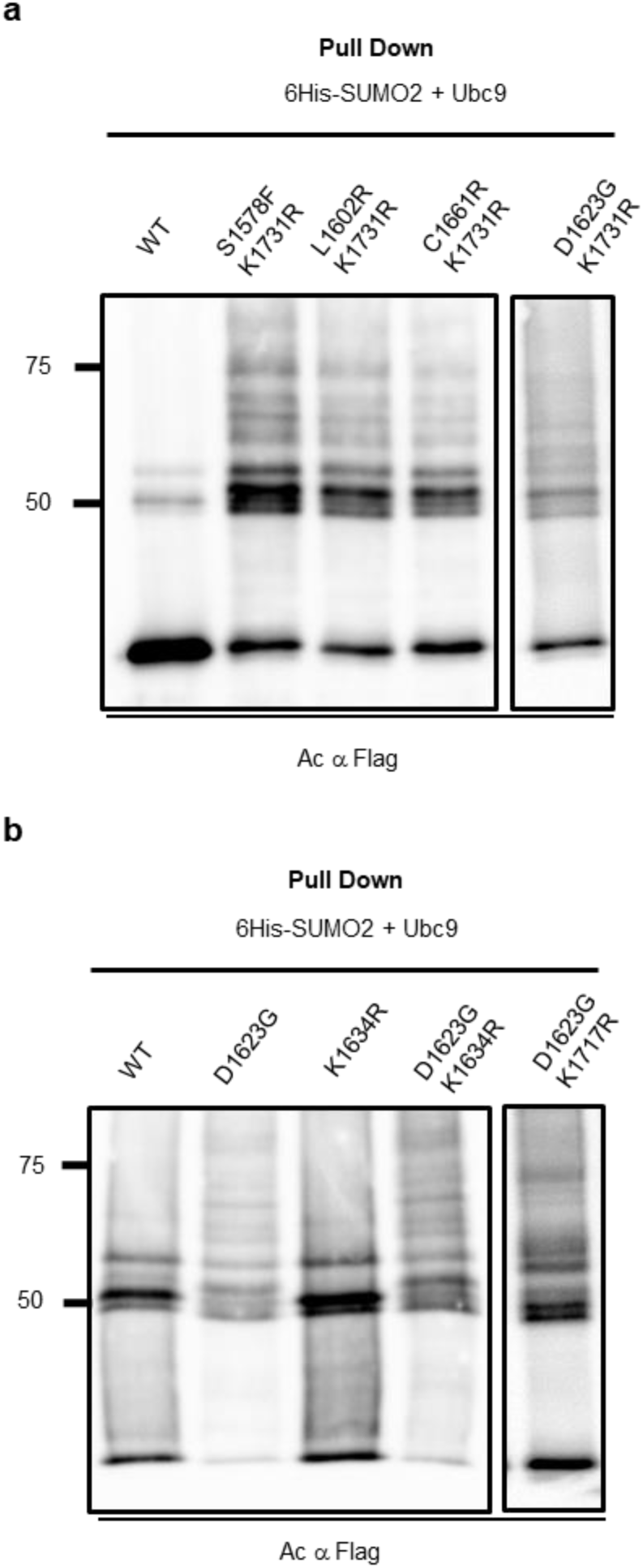
The specific SUMOylation pattern of various patient mutants does not involve K1634 or K1731 but involves K1717 (related to Fig. 7) HEK293 cells were co-transfected with pcDNA3 6His-SUMO2, pcDNA3 Ubc9, and plasmids encoding in (**a**): p3XFlag SecPH, or p3XFlag SecPH derivatives harbouring the KR substitution at the major sumoylation site K1731 and either of the following patient missense mutations S1578F, L1602R, D1623G, and C1661R; and in (**b**): p3XFlag SecPH double mutants harbouring the patient missense mutation D1623G, and an individually mutated lysine belonging to predicted SUMOylation consensus sites **K1634R**, or **K1717R**. Cell lysates were subjected to pull-down assays and analysed as in Fig. 2c. The different panels in (**a**) and (**b**) show different exposure times. The nonspecific binding of unmodified SecPH to the cobalt beads is indicated by an asterisk.

**Supplementary Figure 6.**
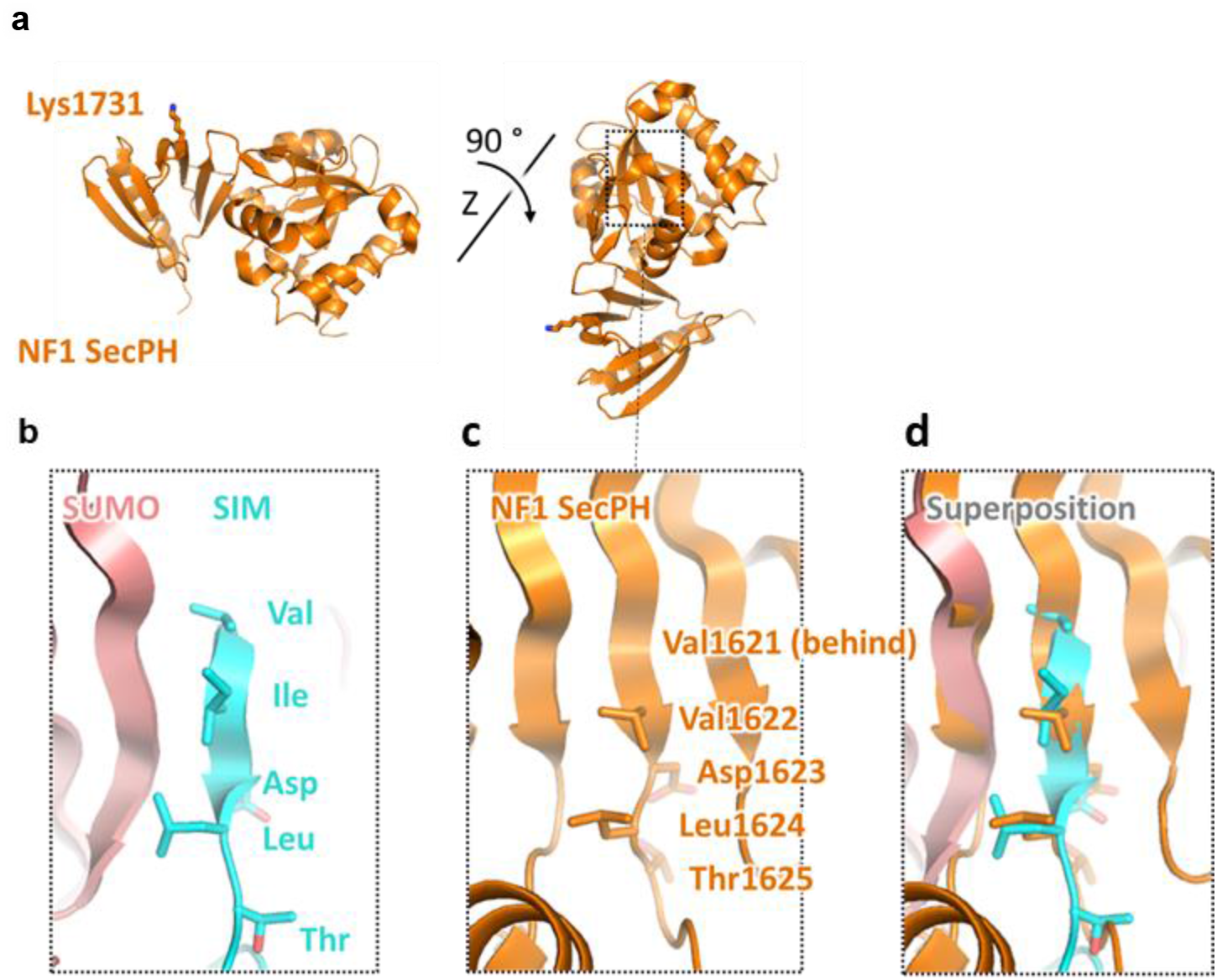
A predicted SecPH SIM domain which could be involved in K1731 SUMOylation is not accessible. **a** SecPH (PDB: 2E2X) is shown in the same orientation as in Fig. 5 and after the indicated rotation. A fragment zoomed in in (**c**) is marked with a dashed square. The side chain of the main SUMOylation site, K1731, is shown as red sticks. **b** The molecular interface between the SIM^CAF1^ and SUMO3 (PDB: 2RPQ). SUMO is shown in ribbon representation in pink and SIM^CAF1^ is shown in stick representation in blue. **c** A fragment of the Nf1 SecPH domain 3D structure surrounding its predicted SIM motif located in the central β-sheet of the Sec domain. Residues corresponding to this predicted SIM are labelled and their side chains are shown. **d** The superposition of (**b**) and (**c**) demonstrates that the predicted SIM domain in SecPH, due to being located on the central β-strand of a β-sheet, is unavailable for binding to SUMO.

**Supplementary Figure 7.**
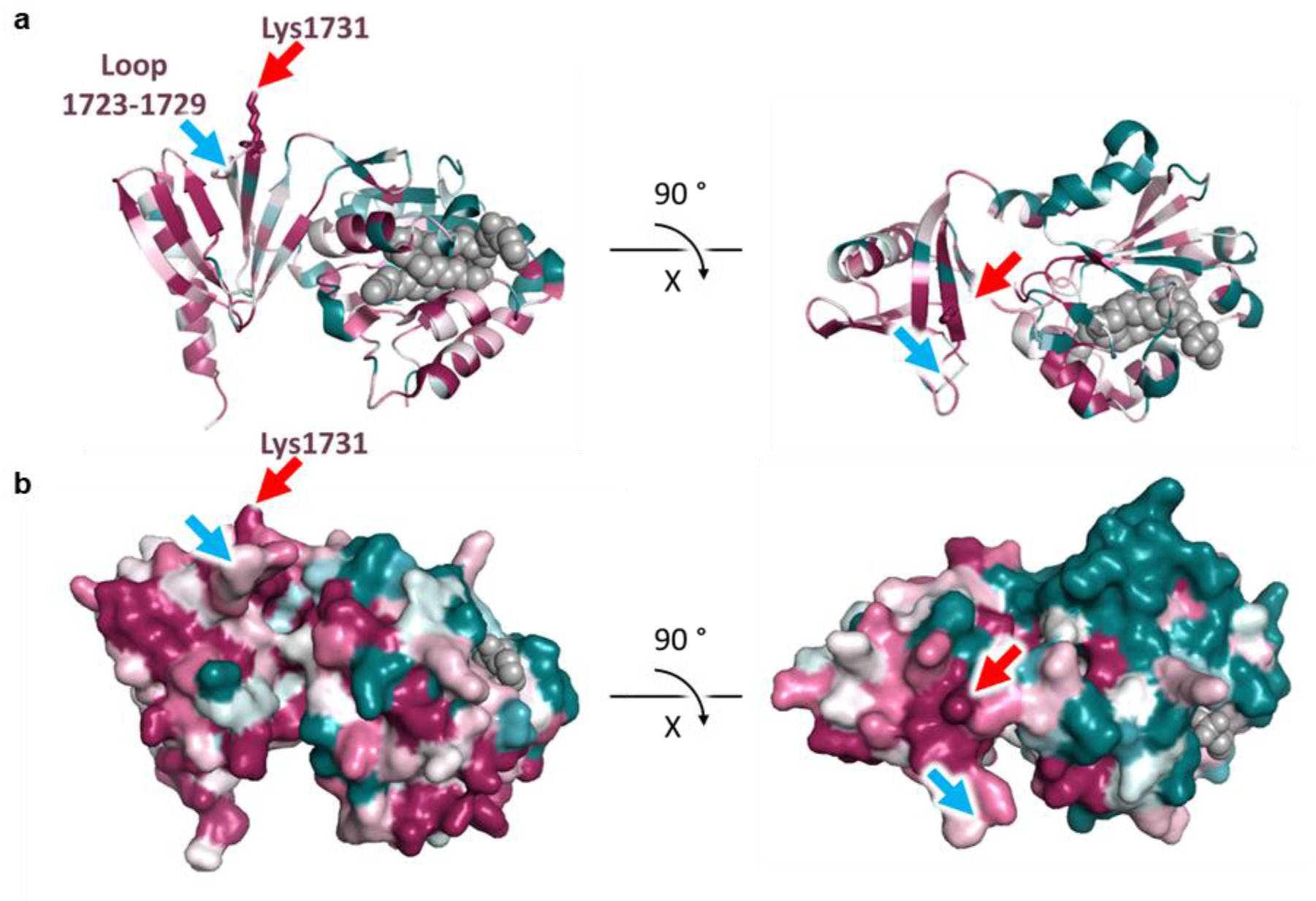
K1731 is highly conserved in most metazoan Nf1 orthologues. **a** SecPH sequence was analysed using ConSurf software as described in Methods and coloured according to sequence conservation. SecPH (PDB: 2E2X) is shown in ribbon representation in the same orientation as in Fig. 5 and after the indicated rotation. The side chain of the main SUMOylation site, K1731, is shown as sticks and indicated with a red arrow. The neighbouring loop between residues L1723 and D1729 is indicated with a blue arrow. Phosphatidylethanolamine is shown as grey spheres. **b** The same views as in (**a**) but in surface representation.

**Supplementary Figure 8.**
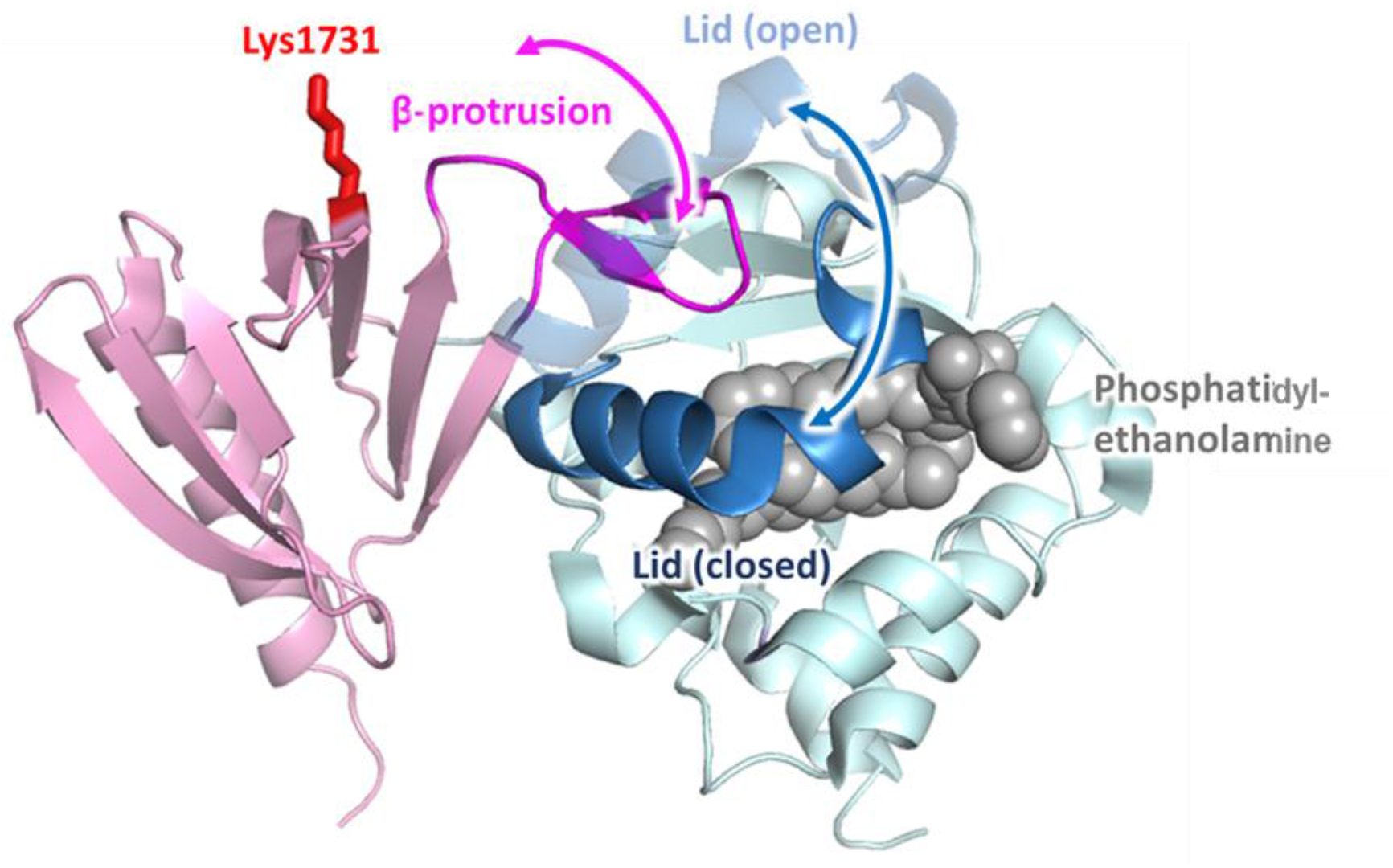
Crucial position of K1731 in the proposed structural rearrangement of the PH β-protrusion and Sec lid dynamics. Ribbon representation of the neurofibromin-SecPH 3D structure complexed with phosphatidylethanolamine (with sphere representation) (PDB: 2E2X), with Sec in light blue, PH in pink, and phosphatidylethanolamine in grey. The K1731 side chain is represented with sticks and labelled in red. The open conformation of the Sec lid is modelled based on a homologous structure of *Saccharomyces cerevisiae* phosphatidylinositol-transfer protein Sec14p (PDB: 1AUA).

**Supplementary Table 1:**
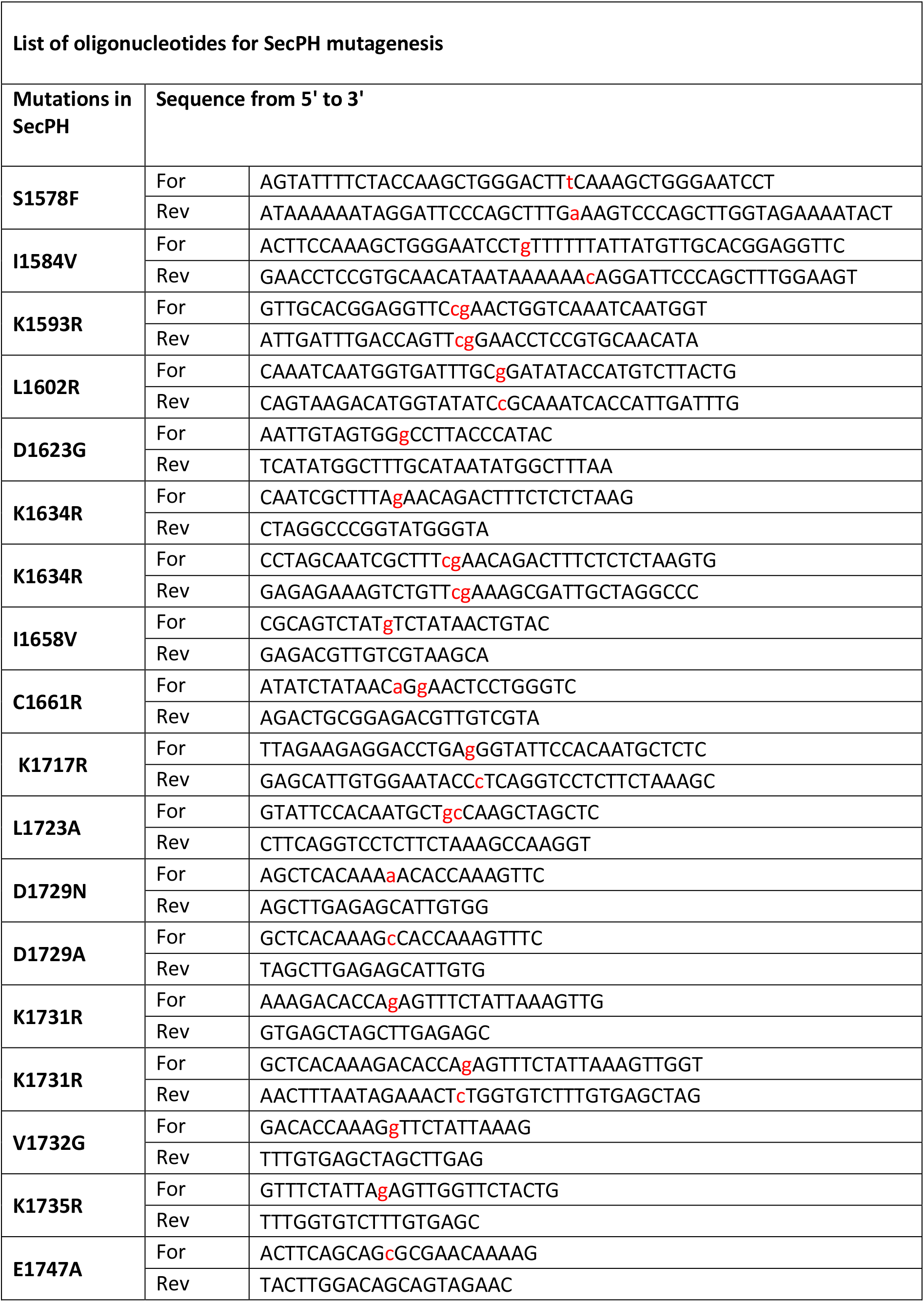
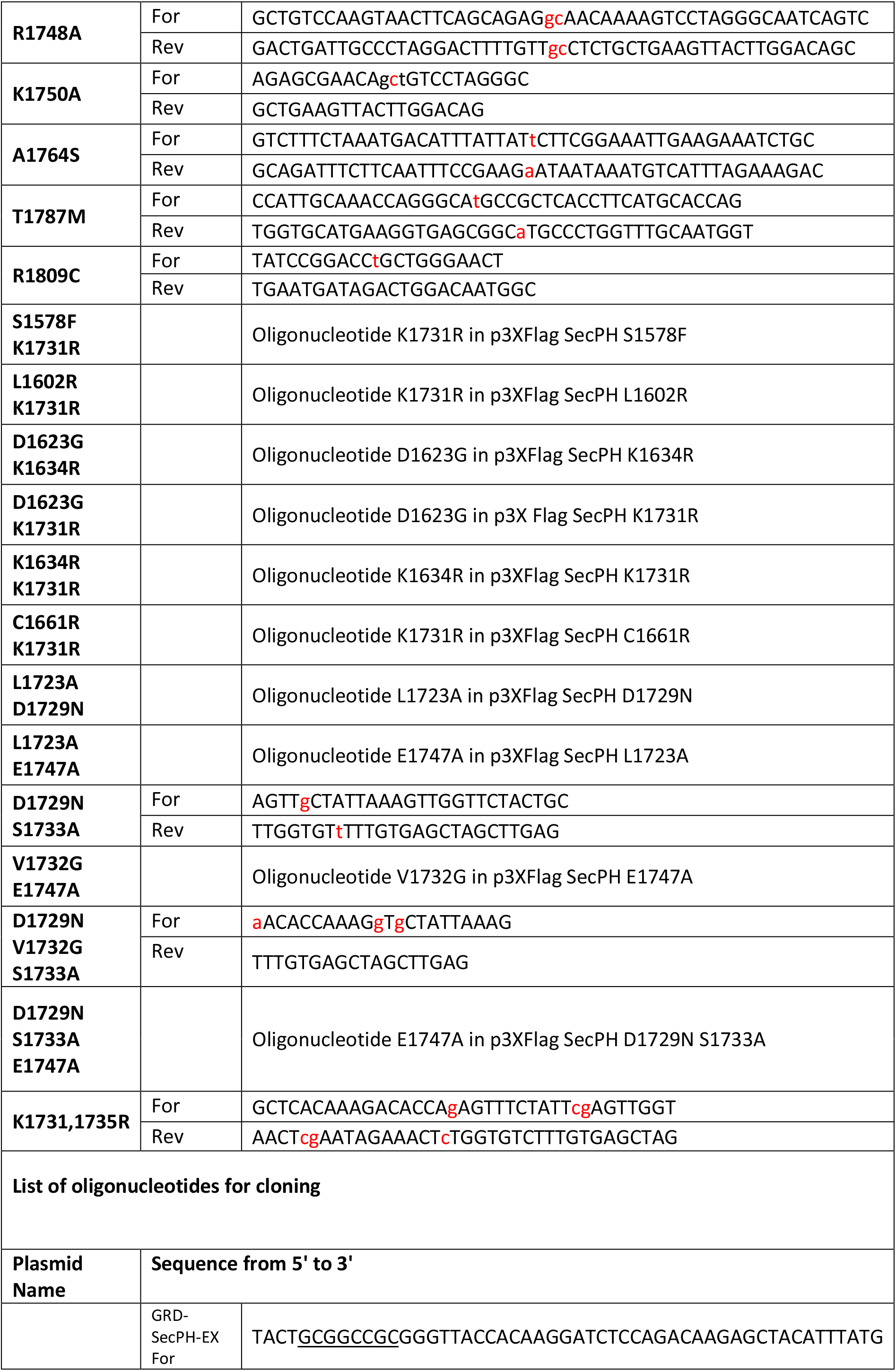
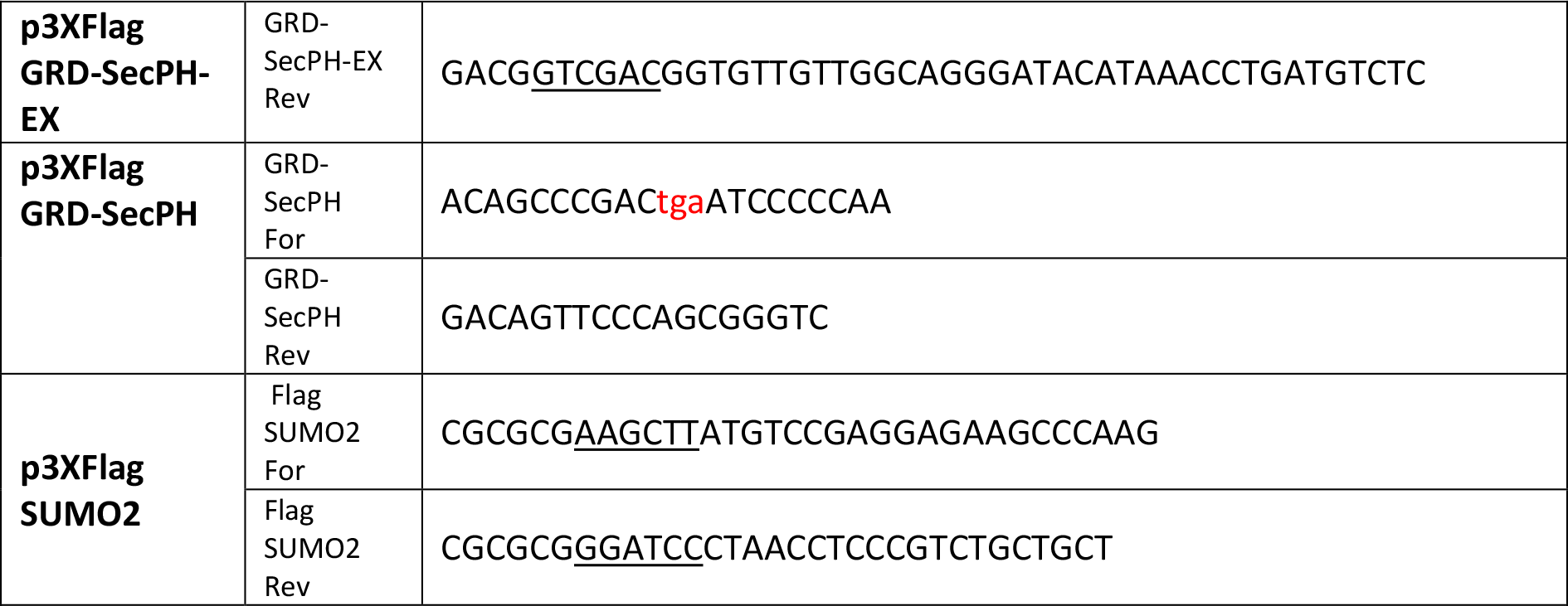
List of oligonucleotides. Mutations introduced are in red and restriction sites are underlined.

**Supplementary Table 2:** List of primary antibodies.

| Primary antibodies |  |  |  |  |
| --- | --- | --- | --- | --- |
|  | Dilution | Biological source | Company | Reference |
| Flag | 1:5000 | Mouse | Sigma-Aldrich | F3165 |
| SUMO2/3 | 1:2000 | Mouse | Abcam | ab81371 |
| Nf1 (sc-67) | 1:5000 for immunodetection of endogeneous Nf1 in HeLa cells | Rabbit | Santa Cruz Biotechnology, Inc. | sc-67 discontinued |
| Nf1 | Undiluted in endogeneous Nf1 IP experiment in HEK293T cells | Rabbit | Novus Biologicals | NB100-418 |
| P-ERK | 1:2000 | Rabbit | Cell Signaling Technology | #4370 |
| ERK | 1:1000 | Rabbit | Cell Signaling Technology | #4695 |
| $\alpha$ Tubuline | 1:1000 | Mouse | Santa Cruz Biotechnology, Inc. | sc-8065 |

**Supplementary Table 3:**
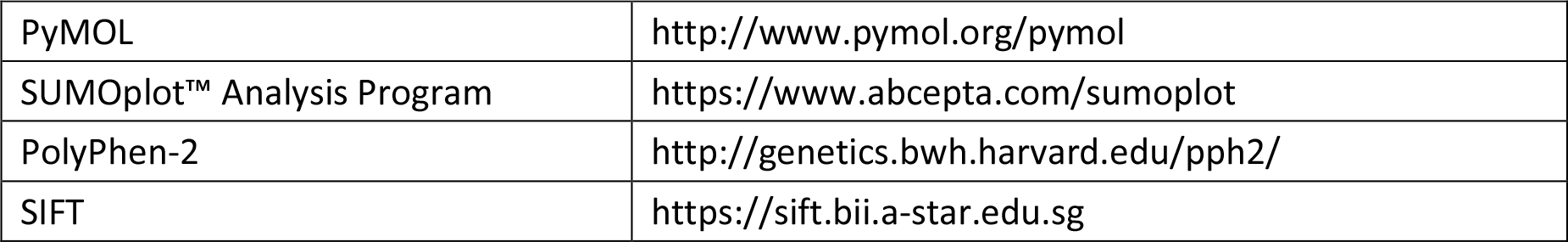
Software.

|  |  |
| --- | --- |
| PyMOL | http://www.pymol.org/pymol |
| SUMOPlot™ Analysis Program | https://www.abcepta.com/sumoplot |
| PolyPhen-2 | http://genetics.bwh.harvard.edu/pph2/ |
| SIFT | https://sift.bii.a-star.edu.sg |

**Supplementary Table 4:** Databases.

|  |  |
| --- | --- |
| ClinVar | https://www.ncbi.nlm.nih.gov/clinvar/ |
| gnomAD | https://gnomad.broadinstitute.org/gene/ENSG00000196712?dataset=gnomad_r2_1 |

**Correspondence** and requests for materials should be addressed to H.B.

